# Enhanced replication of mouse adenovirus type 1 following virus-induced degradation of protein kinase R (PKR)

**DOI:** 10.1101/584680

**Authors:** Danielle E. Goodman, Carla D. Pretto, Tomas A. Krepostman, Kelly E. Carnahan, Katherine R. Spindler

## Abstract

Protein kinase R (PKR) plays a major role in activating host immunity during infection by sensing dsRNA produced by viruses. Once activated by dsRNA, PKR phosphorylates the translation factor eIF2α, halting cellular translation. Many viruses have methods of inhibiting PKR activation or its downstream effects, circumventing protein synthesis shutdown. These include sequestering dsRNA or producing proteins that bind to and inhibit PKR activation. Here we describe our finding that in multiple cell types, PKR was depleted during mouse adenovirus type 1 (MAV-1) infection. MAV-1 did not appear to be targeting PKR at a transcriptional or translational level because total PKR mRNA levels and levels of PKR mRNA bound to polysomes were unchanged or increased during MAV-1 infection. However, inhibiting the proteasome reduced the PKR depletion seen in MAV-1-infected cells, whereas inhibiting the lysosome had no effect. This suggests that proteasomal degradation alone is responsible for PKR degradation during MAV-1 infection. Time course experiments indicate that the degradation occurs early after infection. Infecting cells with UV-inactivated virus prevented PKR degradation, whereas inhibiting viral DNA replication did not. Together these results suggest that an early viral gene is responsible. Degradation of PKR is a rare mechanism to oppose PKR activity, and it has only been described in six RNA viruses. To our knowledge, this is the first example of a DNA virus counteracting PKR by degrading it.

**Importance:** The first line of defense in cells during viral infection is the innate immune system, which is activated by different viral products. PKR is a part of this innate immune system and is induced by interferon and activated by dsRNA produced by DNA and RNA viruses. PKR is such an important part of the antiviral response that many viral families have gene products to counteract its activation or the resulting effects of its activity. Although a few RNA viruses degrade PKR, this method of counteracting PKR has not been reported for any DNA viruses. MAV-1 does not encode virus-associated RNAs, a human adenoviral defense against PKR activation. Instead, MAV-1 degrades PKR, and it is the first DNA virus reported to do so. The innate immune evasion by PKR degradation is a previously unidentified way for a DNA virus to circumvent the host antiviral response.

## Introduction

Activation of protein kinase R (PKR) is a major innate immune response to viral infection. PKR is an interferon (IFN)-induced protein that is comprised of two major domains, an N-terminal double-stranded RNA binding domain and a C-terminal serine/threonine kinase domain (1, 2). PKR binds to dsRNA (3-5) and once bound, it becomes activated by dimerizing and autophosphorylating (6-9). When activated, PKR phosphorylates eukaryotic translation initiation factor 2α (eIF2α), causing inhibition of protein synthesis and reduced viral replication (10-13). Many viruses encode gene products that block PKR activation or inhibit its ability to phosphorylate eIF2α (14). A common mechanism is to produce a viral protein that binds and sequesters dsRNA, blocking its interaction with PKR. Examples of this are vaccinia virus E3L (15-17), influenza virus NS1 (18, 19), and Ebola virus protein VP35 (20). Other viruses produce proteins or RNA that bind directly to PKR to inhibit its activation, such as herpes simplex virus US11 (21, 22), HIV-1 Tat protein (23, 24) or TAR RNA (25), and human adenovirus (hAd) virus-associated (VA) RNAs (10, 26-28).

Degradation of PKR by viruses is a less documented method of regulating PKR. To date PKR degradation has been reported in six RNA viruses: Toscana virus (TOSV) (29), Rift Valley fever virus (RVFV) (30-32), poliovirus (33, 34), foot-and-mouth disease virus (FMDV) (35, 36), encephalomyocarditis virus (EMCV, strain mengovirus) (37, 38), and enterovirus 71 (39). RVFV and TOSV both degrade PKR via proteasomal mechanisms involving a viral nonstructural protein (NSs) (32, 40, 41). RVFV NSs recruits a SCF (SKP1-CUL1-F-box)^FBXW11^ E3 ubiquitin ligase to ubiquitinate PKR and target it to the proteasome, though PKR ubiquitination could not be demonstrated (32, 41). The mechanism for PKR proteasomal degradation by NSs has not been described for TOSV (40). FMDV uses the other major cellular protein degradation pathway, the lysosome, to degrade PKR during infection (36). Though the mechanism is unclear, expression of the major FMDV protease 3C^pro^ is required for PKR degradation by the lysosome. However, 3C^pro^ does not interact with PKR, nor is its protease activity required for PKR degradation. The enterovirus A71 3C^pro^ causes PKR degradation by direct interaction (39). The mechanism of PKR depletion by poliovirus is unclear, though gene expression is required, and the major poliovirus proteases (2A and 3C) are not directly involved (33). The mechanism by which mengovirus depletes PKR during infection is unknown (37, 38).

Adenoviruses are species-specific, making the study of hAd pathogenesis difficult in an animal model. MAV-1 is a useful alternative to study adenovirus pathogenesis (42-46). MAV-1 has molecular, genetic, and pathogenic similarities and differences to hAd. Their genomic structures are similar at a gross level, and both contain early genes involved in pathogenesis and immune evasion. Pathogenically, their tropisms vary, with hAd infecting epithelial cells, leading to upper respiratory and GI tract infections, and conjunctivitis, while MAV-1 infects endothelial cells and monocytes, causing encephalitis and myocarditis. We and others have been investigating the adaptive and innate immune responses to MAV-1.

Human adenovirus VA RNAs bind PKR as a monomer, preventing its transautophosphorylation (47). However, MAV-1 does not produce VA RNAs (48), and it is not known whether MAV-1 induces PKR activation. In our studies of MAV-1 pathogenesis and the innate response, we discovered that during MAV-1 infection, PKR was depleted from cells as early as 12 hours post infection (hpi). Total PKR mRNA levels and PKR mRNA bound to polysomes were unchanged or increased during MAV-1 infection, suggesting that MAV-1 did not appear to be targeting PKR at a transcriptional or translational level. However, inhibiting the proteasome blocked the PKR depletion seen in MAV-1-infected cells, indicating that proteasomal degradation is responsible for PKR depletion during MAV-1 infection. We report results indicating that an early viral gene is likely responsible for mediating PKR degradation. To our knowledge, this is the first example of a DNA virus counteracting PKR by degrading it.

## Results

### Viral DNA yield is increased in PKR^−/−^ mouse embryonic fibroblasts

While PKR is an important part of the innate immune response, PKR^−/−^ cells in culture are not always more susceptible to viral infection than wild type cells (49-51). PKR^−/−^ mouse embryonic fibroblasts (MEFs) show increased viral yields compared to wild type MEFs when infected with vesicular stomatitis virus and influenza A (49, 50), but there is no change in viral yield during vaccinia virus infection compared to wild type cells (51). However, it was later discovered that the PKR^−/−^ MEF lines used are not complete PKR knockouts (52). There are two categories of PKR^−/−^ MEFs derived from knockout mice: N-PKR^−/−^ MEFs and C-PKR^−/−^ MEFs (52). The PKR^−/−^ MEFs derived from mice created in the Weissmann lab (53) are designated N-PKR^−/−^ MEFs, because the C-terminal fragment of PKR is still expressed and can be detected by immunoblot when there is IFN induction (52). The fragment has the kinase catalytic activity of PKR, but it does not bind dsRNA (52). The PKR^−/−^ MEFs derived from mice created in the Bell lab (54) are designated as C-PKR^−/−^ MEFs, because the N-terminal fragment of PKR is still expressed and can be detected by immunoblot with specific PKR antibodies (52). The fragment is catalytically inactive, but it can still bind dsRNA (52). Susceptibility of these PKR^−/−^ MEFs to specific viruses may be dependent on the PKR mutation and the mechanism used by each virus to circumvent PKR.

To determine whether PKR plays an important role during MAV-1 infection, we tested the susceptibility of both PKR^−/−^ MEF lines to MAV-1 infection. We infected wild type MEFs, N-PKR^−/−^ MEFs, and C-PKR^−/−^ MEFs with MAV-1 at an MOI of 1 PFU/cell and collected cell pellets at 48 and 72 hpi. DNA was purified from the cell pellets and analyzed for MAV-1 genome copies by qPCR. N-PKR^−/−^ MEFs produced a significantly higher viral DNA yield than wild type MEFs at 48 hpi, and both PKR mutant MEF lines had a significantly higher viral DNA yield than wild type MEFs at 72 hpi (Fig. 1). Although we have not confirmed the production of truncated PKR proteins in the cells in our laboratory, the results of Fig. 1 indicate that PKR activation is an important antiviral response during MAV-1 infection *in vitro*.

**Figure 1.**
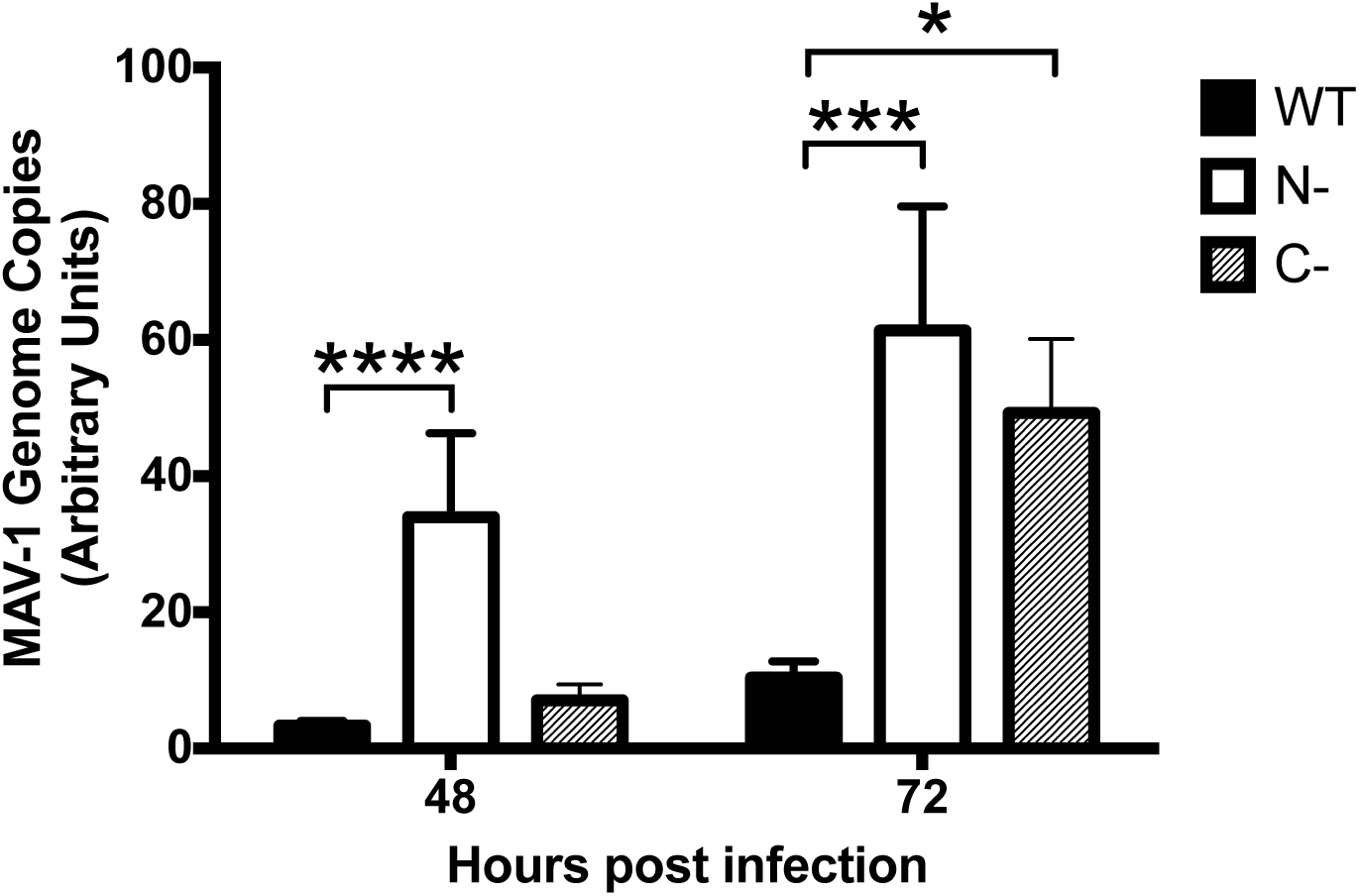
Viral DNA yield is increased in PKR^−/−^ mouse embryonic fibroblasts. PKR WT MEFs (WT), N-PKR^−/−^ MEFs (N-), and C-PKR^−/−^ MEFs (C-) were infected with MAV-1 at an MOI 1 and collected at 48 and 72 hpi. DNA was purified from cell pellets and analyzed for MAV-1 genome copies by qPCR. Graph is representative of three experiments, 14 biological replicates per cell line per time point. Error bars are standard error of the mean (SEM). \**P* ≤ 0.05, \*\*\**P* ≤ 0.0002, \*\*\*\**P* ≤ 0.0001.

### Mouse PKR is depleted during MAV-1 infection

To determine whether MAV-1 affects PKR during infection, we infected several cell types and analyzed PKR protein expression. We infected immortalized C57BL/6 MEFs, C57BL/6 primary peritoneal macrophages, and CMT93 cells (mouse rectal carcinoma cells) with MAV-1 at an MOI of 10 and collected cell lysates 24, 48, and 72 hpi. We analyzed cell lysates for the presence of PKR by immunoblot using a polyclonal antibody that detects mouse PKR. We probed blots with antibodies to actin as a loading control. To our surprise, in C57BL/6 MEFs, PKR was almost completely depleted from lysates 24 hpi and remained depleted through 72 hpi (Fig. 2A and B). PKR was also depleted compared to mock infection at 24 and 48 hpi in C57BL/6 MEFs infected at MOI 2 and 5 (Fig. S1). We also observed depletion of PKR in other cell types. In CMT93 cells, PKR was nearly undetectable at 24 hpi (Fig. 2A). In C57BL/6 primary peritoneal macrophages, PKR was decreased at 48 hpi compared to mock lysates, and absent in infected lysates at 72 hpi (Fig. 2A). This indicates that MAV-1 causes PKR depletion during infection.

**Figure 2.**
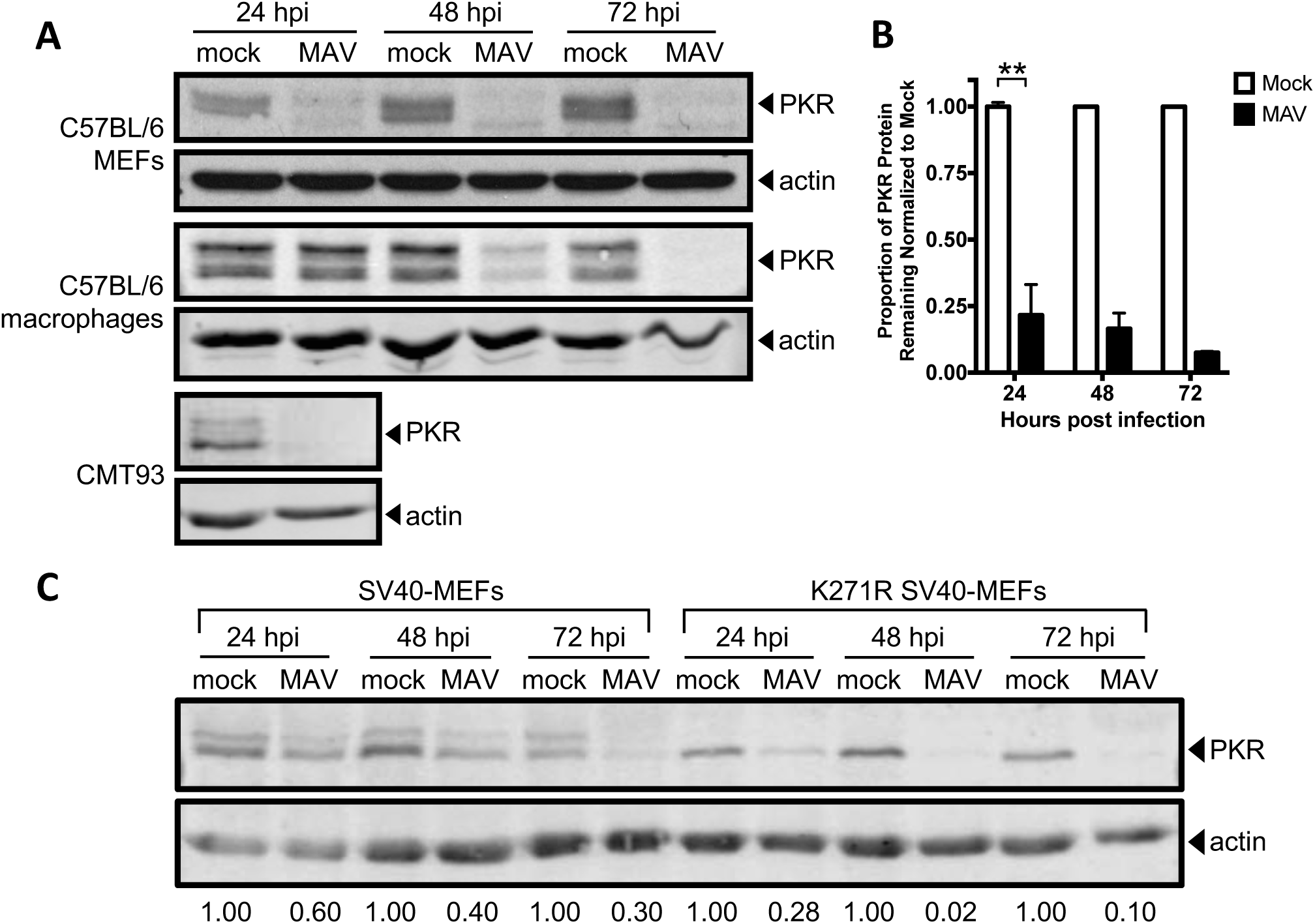
Mouse PKR is depleted during MAV-1 infection. (A) Cells (indicated at left) were infected with MAV-1 (MAV) at an MOI of 10 or mock infected (mock). Cell lysates were collected at the indicated times and analyzed by immunoblot with antibodies for PKR (B-10 for C57BL/6 MEFs and D-20 for C57BL/6 primary peritoneal macrophages and CMT93 cells) and actin. Blots are representative of a minimum of three independent experiments per cell line. (B) Densitometry quantitation of five C57BL/6 MEFs immunoblots from A. Error bars are SEM. N=5 for 24 hpi, n=4 for 48 hpi, and n=2 for 72 hpi. \*\**P* ≤ 0.01. (C) SV40-MEFs or kinase-dead (K271R) SV40-MEFs were infected with MAV-1 at an MOI of 10. Cell lysates were immunoblotted as in A with PKR D-20. Numbers below are the proportion of PKR protein for each time point, normalized to actin and the mock PKR protein levels from the corresponding time point.

To determine whether kinase activity of PKR is important for the depletion, we assayed infection of MEFs expressing a mutant form of mouse PKR with a point mutation in the kinase domain (K271R) (55). These cells, designated K271R SV40-MEFs, showed an even more rapid depletion of PKR than in WT SV40-MEFs (Fig. 2C). At 24 hpi, in K271R SV40-MEFs, 28% of PKR remained, compared to 60% in the WT SV40-MEFs. The fraction remaining at 72 hpi in K271R SV40-MEFs was 10%, compared to 30% in WT SV40-MEFs (Fig. 2C). This indicates that the PKR kinase does not have to be functional to be depleted during MAV-1 infection. Also, comparing the PKR immunoblot bands in the mock-infected WT SV40 MEFs and mutant K271R SV40-MEFs suggests that the upper band of the PKR doublet usually seen in wild-type cells is a phospho-PKR band, because only the lower band of the PKR doublet is seen in kinase-dead mutant K271R SV40-MEFs. The data in Fig. 2A and C thus indicate that both PKR and phospho-PKR are depleted during MAV-1 infection.

### MAV-1 does not cause PKR depletion by reducing steady-state levels of PKR mRNA

To determine the mechanism of PKR depletion, we first assayed whether reduction in PKR protein during MAV-1 infection was due to reduced PKR mRNA steady-state levels. We mock-infected or infected C57BL/6 MEFs and primary peritoneal macrophages at an MOI of 10 and collected cell lysates at 24, 48, and 72 hpi. We synthesized cDNA from RNA purified from these cell lysates and assayed for PKR mRNA by qPCR. In C57BL/6 MEFs, PKR mRNA levels were similar between mock and infected lysates at 24 hpi (Fig. 3A), a time point in which PKR protein levels were already greatly depleted in the infected lysates compared to mock lysates (Fig. 2). Although PKR mRNA levels were depleted 33% at 48 hpi and 40% 72 hpi in MAV-1-infected lysates compared to mock lysates, this does not correlate to the 84% and 94% reduction, respectively, in PKR protein levels at those time points (Fig. 2B). In C57BL/6 primary peritoneal macrophages, PKR mRNA levels in the infected lysates were 2-3 times higher than levels in mock lysates at all three time points assayed (Fig. 3B), even though PKR protein levels were almost completely depleted in infected lysates at 72 hpi (Fig. 2). This is evidence that MAV-1 is not causing PKR protein depletion by reducing PKR steady-state mRNA levels during infection.

**Figure 3.**
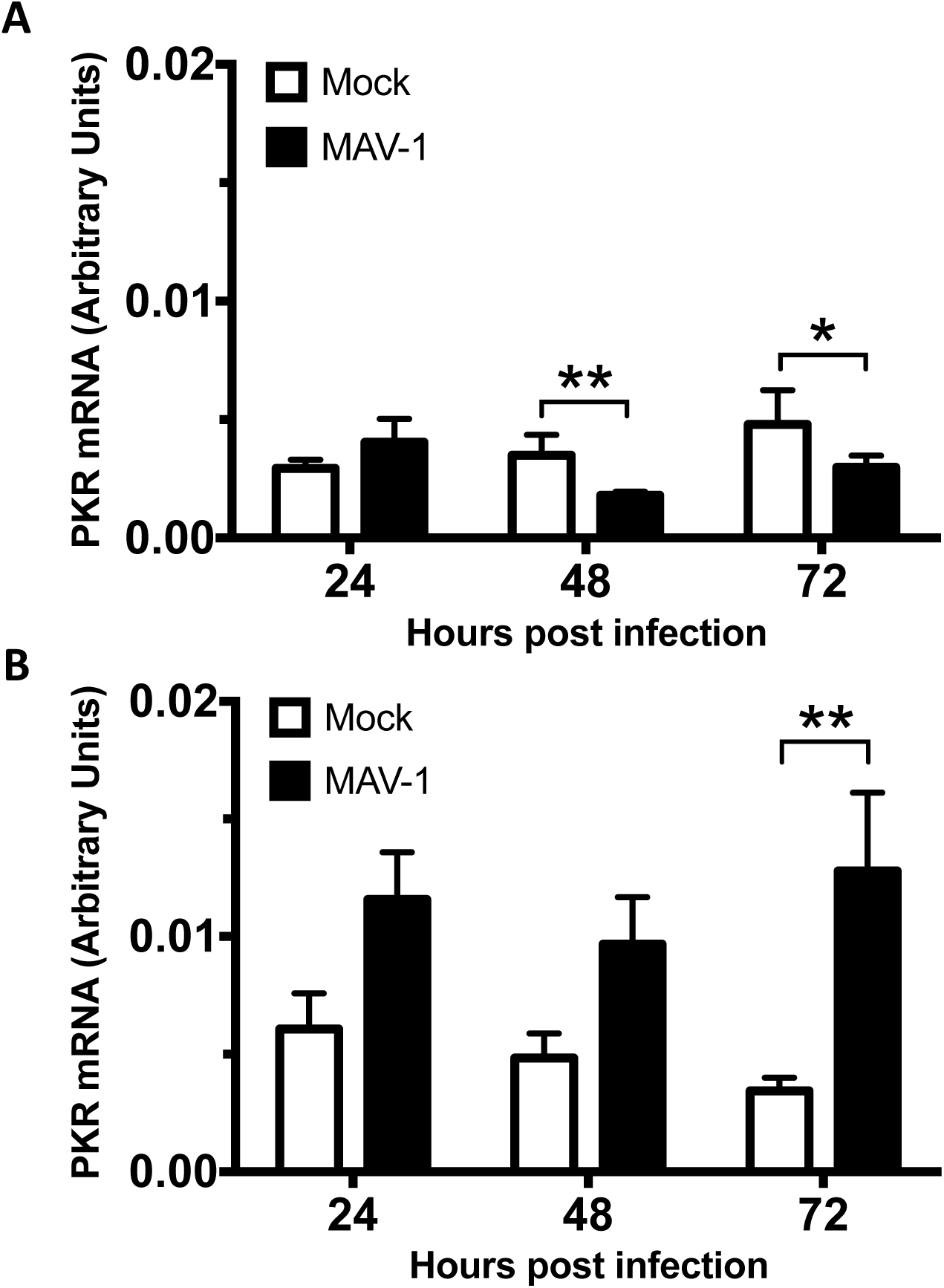
MAV-1 does not cause PKR depletion by reducing steady state levels of PKR mRNA at times when the protein levels are already reduced. MEFs (A) or isolated primary peritoneal macrophages (B) were harvested and infected with MAV-1 at an MOI of 10 or mock infected. The cell pellets were collected and RNA was isolated. cDNA was generated from the RNA, and qPCR was used to quantitate PKR mRNA levels. Each graph contains 5-7 replicates for each time point from three pooled experiments. Error bars show the SEM. \**P* ≤ 0.05 and \*\**P* ≤ 0.01.

### MAV-1 infection effects on PKR translation

Because MAV-1 did not reduce PKR mRNA steady-state levels, we determined whether MAV-1 causes PKR depletion by reducing translation of its mRNA. We first assayed total PKR mRNA bound to ribosomes during infection. C57BL/6 MEFs were mock infected or infected at an MOI of 5, and lysates were collected at 48 hpi in the presence of cycloheximide to keep the mRNA bound to the ribosomes (56). Lysates were centrifuged through 25% sucrose to pellet ribosomes, and RNA was purified from the pellets. The purified RNAs were used to generate cDNA, which we assayed for PKR mRNA by qPCR. As a control for pelleting of ribosomes, we assayed the pellets and sucrose cushion supernatants by immunoblot with antibodies to ribosomal protein RPL7. We confirmed that RPL7 was only present in the pellets and not the supernatants (Fig. S2). There was no significant difference between the amount of PKR mRNA in the ribosome pellet of mock infected lysates compared to MAV-1-infected lysates (Fig. 4A).

**Figure 4.**
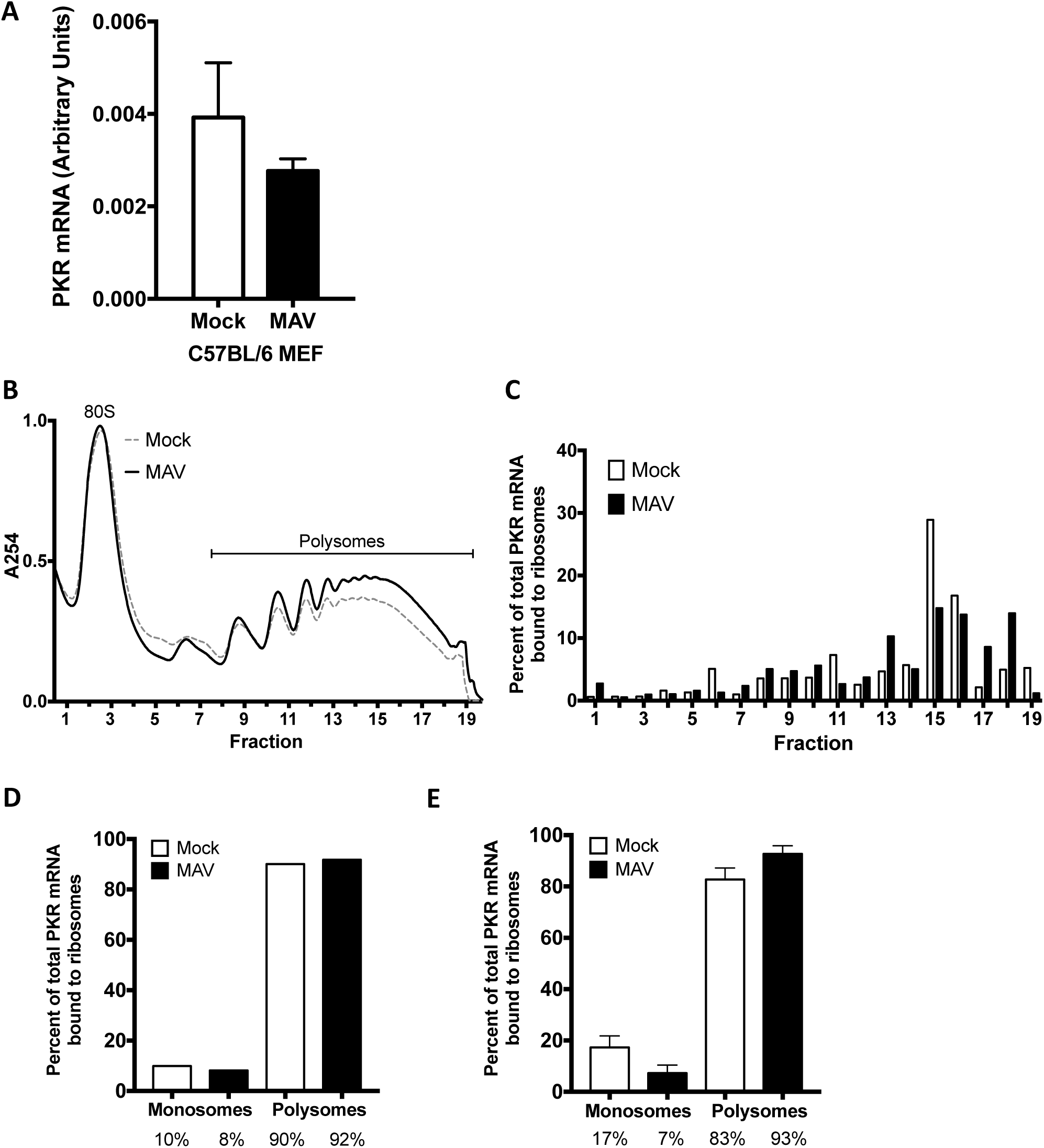
MAV-1 infection does not affect PKR translation. (A) C57BL/6 MEFs were infected with MAV-1 (MAV) at an MOI of 5 or mock infected (mock) and collected at 48 hpi. Cells were lysed, and cleared lysates from three 10 cm plates were layered onto 25% sucrose and centrifuged to pellet ribosomes. RNA was purified from the pellets, cDNA was generated from the RNA, and qPCR was used to quantitate the PKR mRNA levels. The graph contains 9 replicates for each time point, pooled from 3 independent experiments. Error bars show the SEM. (B) C57BL/6 MEFs were infected with MAV-1 (MAV) at an MOI of 2 or mock infected (mock). Cells were collected at 25 hpi and lysed; cleared lysates were layered onto 10-50% sucrose gradients and centrifuged. Gradients were collected from the top and pumped through a UV spectrophotometer, and 34 fractions were collected. The gradients are displayed with the bottom fractions to the right. The UV trace of the first 10 fractions (including 40S and 60S ribosomal subunits) is not shown. (C) RNA was purified from each fraction of the gradients in B. cDNA was generated from the RNA, and qPCR was used to quantitate PKR mRNA in each fraction and displayed as the percent of total PKR mRNA associated with ribosomes. B and C are results from one representative experiment of 3 independent experiments. (D) Percentage of total PKR mRNA associated with monosomes (fractions 1-6) and polysomes (fractions 7-19) from the trial displayed in B and C were pooled for mock and infected samples. The percentages represented by each bar are displayed below each bar. (E) Pooled monosome and polysome data as described in D from three independent experiments. Error bars show the SEM. The percentages represented by each bar are displayed below each bar. There were no significant differences between mock and infected samples (A, E).

To confirm the results seen in total mRNA bound to ribosomes, we also centrifuged cell extracts on sucrose gradients to generate polysome profiles. This enabled us to analyze levels of PKR mRNA associated with actively-translating ribosomes during infection. C57BL/6 MEFs were mock infected or infected at an MOI of 2, and lysates were collected at 24 hpi in the presence of cycloheximide, as above. RNA content for mock and infected lysates was estimated by NanoDrop spectrophotometry, and equivalent OD amounts of RNA were centrifuged on 10-50% sucrose gradients to sediment 40S and 60S ribosomal subunits, 80S ribosomes (monosomes), and polyribosomes (polysomes). A typical polysome profile was obtained (Fig. 4B). RNA was purified from fractions containing monosomes and polysomes and then used to generate cDNA, which we assayed for PKR mRNA by qPCR (Fig. 4C). As a control, GAPDH mRNA was measured by qPCR, and PKR mRNA levels in each fraction were normalized to the GAPDH mRNA content. When the data for percentage of PKR mRNA bound to ribosomes was pooled into monosome and polysome fractions and analyzed (Fig. 4D), 90.1% and 91.8% were bound to polysomes (fractions 7-19) for mock and infected samples, respectively, compared to 9.9% and 8.2% bound to monosomes (fractions 1-6). We performed two additional polysome gradient analyses. The pooled data from all three (Fig. 4E) were similar to the data for Fig. 4D, i.e., 82.7% and 92.7% of PKR mRNA were bound to mock and infected polysomes, respectively, compared to 17.3% and 7.3% bound to monosomes. Thus, PKR protein depletion during MAV-1 infection does not appear to stem from a decrease in PKR mRNA translation.

We also assayed whether PKR mRNA might have a signal that would reduce its translation during MAV-1 infection. We constructed a plasmid that positioned sequence corresponding to the 5’ UTR of PKR mRNA upstream of a reporter nanoluciferase gene (57), transfected it into C57BL/6 MEFs, then infected with MAV-1. Compared to cells transfected with a control plasmid with the human β-globin 5’ UTR upstream of the reporter nanoluciferase, there was no significant difference in luciferase activity between mock infected and infected samples (Fig. S3). These data suggest that MAV-1 is not affecting PKR translation through interaction with the 5’ UTR of PKR. The data are consistent with the ribosome pellet and polysome data that MAV-1 infection does not reduce PKR mRNA translation.

### PKR is depleted by proteasomal degradation during MAV-1 infection

There are two main proteolysis pathways in cells, proteasomal degradation and lysosomal degradation (58). To determine whether MAV-1 depletes PKR by either protein degradation pathway, we first assayed whether PKR is lysosomally degraded as follows. CMT93 cells were mock infected or infected with MAV-1 and treated at the time of infection with the lysosome inhibitors ammonium chloride or chloroquine, or water (as a control). At 24 hpi, we collected lysates and analyzed them by immunoblot with antibodies to PKR. In the presence of the lysosomal degradation inhibitors, PKR was depleted by 24 hpi (Fig. 5A), indicating that lysosomal degradation was not the cause of PKR depletion during MAV-1 infection. We confirmed that the inhibitor treatment did block lysosomal degradation by incubating cells with dye-quenched bovine serum albumin (DQ BSA) in addition to the lysosomal inhibitors. DQ BSA is self-quenched until it is digested in the lysosome (59, 60), and imaging confirmed that cells treated with lysosome inhibitors did not fluoresce, but cells treated with the vehicle control (H_2_O) did, as expected (Fig. 5B).

**Figure 5.**
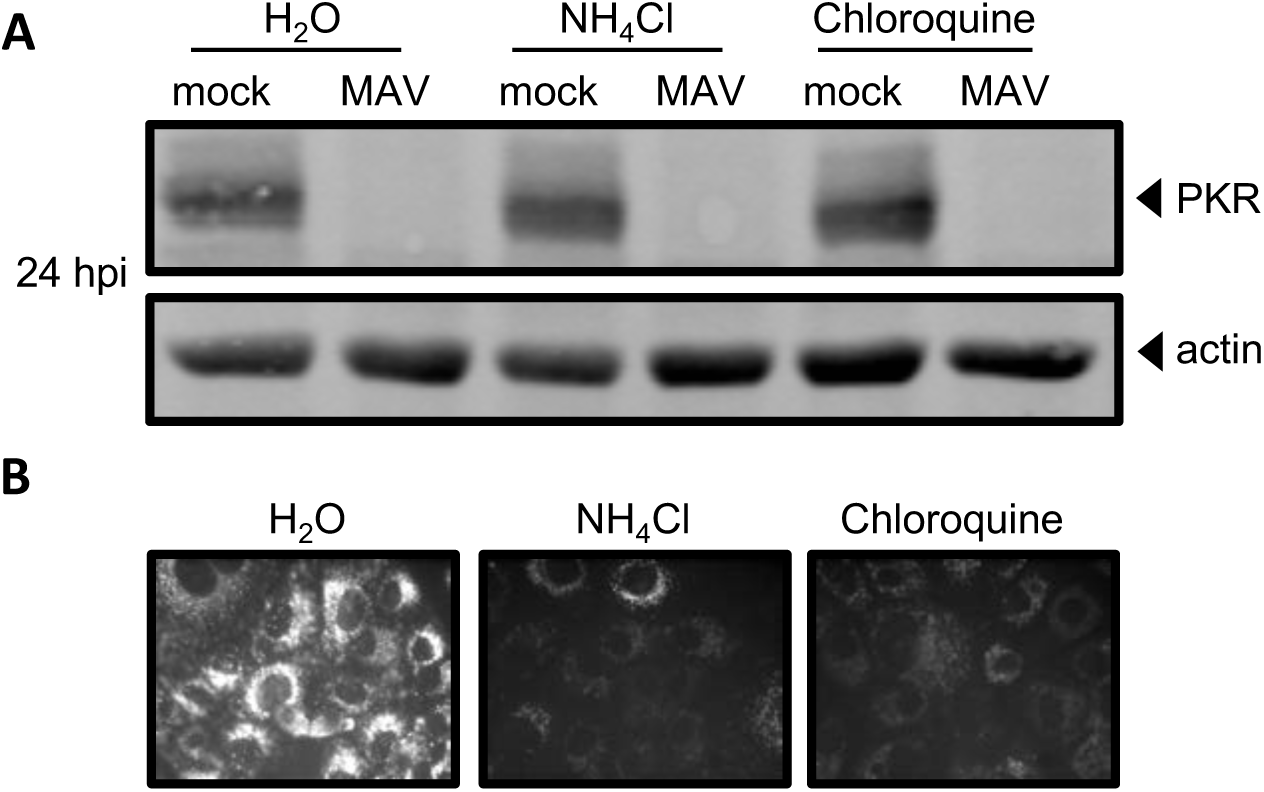
PKR is not depleted by lysosomal degradation during MAV-1 infection. (A) CMT93 cells were infected with MAV-1 (MAV) at an MOI of 10 or mock infected (mock) and treated with 10 mM ammonium chloride or 60 µM chloroquine to inhibit lysosomal degradation, or water, as a control. Cell lysates were analyzed by immunoblot with antibodies for PKR (D-20) and actin. Blots are representative of three independent experiments. (B) Inhibitors were tested for activity using a DQ BSA assay; the DQ BSA molecule will only fluoresce if lysosomal degradation is functional. Uninfected cells were treated as indicated and imaged by fluorescence microscopy.

Next, we examined whether proteasomal degradation is responsible for the degradation of PKR by using proteasome inhibitors MG132 and bortezomib. These inhibit proteasome activity by binding to the active sites in the 20S subunit and blocking the proteolytic activity (61-63). We mock infected or infected C57BL/6 MEFs with MAV-1 and treated with MG132 or bortezomib in DMSO at the time of infection. At 24 hpi, we collected lysates and analyzed them by immunoblot for PKR protein levels. While PKR was depleted in the control DMSO-treated MAV-1-infected cells as expected, PKR protein was present in the MG132- and bortezomib-treated cells at levels comparable to mock infected cells (Fig. 6A and B). To rule out the possibility that PKR was present (not depleted) because the virus infection itself was inhibited by MG132 or bortezomib, we assayed viral replication of MAV-1 with MG132 and bortezomib treatment by qPCR of viral DNA. Viral replication was equivalent in all three treatment groups (Fig. S4), indicating that the treatments did not affect the ability of the virus to productively infect the cells. Taken together, these data indicate that MAV-1 infection results in PKR depletion by causing PKR to be degraded by the proteasome during infection.

**Figure 6.**
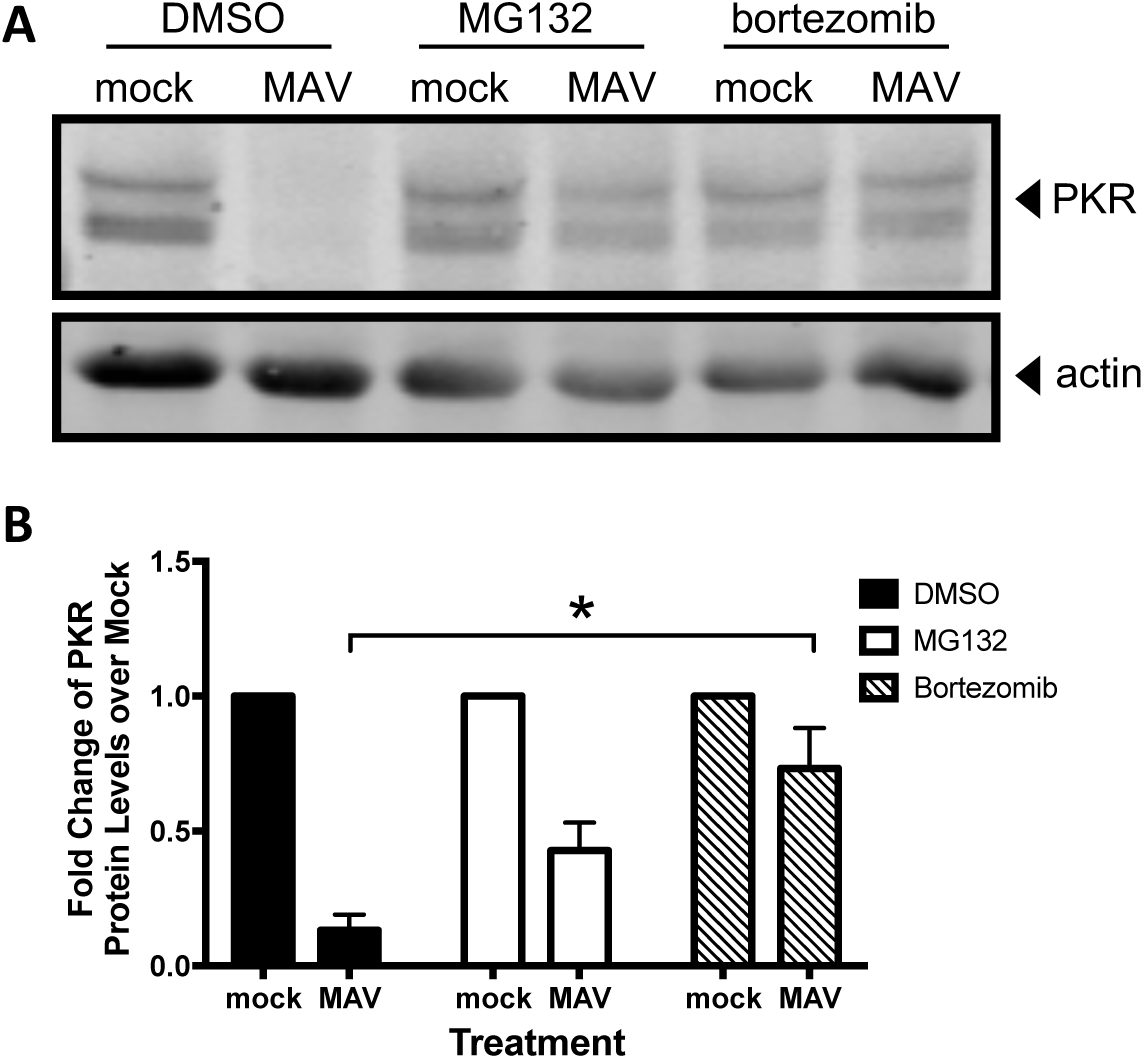
PKR is depleted by proteasomal degradation during MAV-1 infection. (A) C57BL/6 MEFs were infected with MAV-1 (MAV) at an MOI of 10 or mock infected (mock) and treated with DMSO (vehicle for inhibitors), 1 µM MG132, or 1 µM bortezomib. Cell lysates were were analyzed by immunoblot with antibodies for PKR (D-20) and actin. Blots are representative of four independent experiments. (B) Densitometry quantitation of four independent experiments. Treatment with bortezomib significantly inhibited PKR depletion in MAV-1 infected cells, \**P* ≤ 0.05.

A signal for proteasomal degradation is the conjugation of ubiquitin to a protein (64, 65). We examined whether PKR is ubiquitinated by immunoblot. We detected ubiquitination of a positive control, mouse p53, which is degraded in the presence of MAV-1 proteins (66). However, even with the use of epitope-tagged ubiquitin (67) and MG132 treatment, we were unable to detect PKR ubiquitination during infection (Fig. S5). This is consistent with an inability to detect PKR ubiquitination when it is degraded during RVFV infection (32). Although RVFV NSs is known to recruit an E3 ligase to PKR, the authors reported that ubiquitinated PKR is undetectable. Therefore, the cellular degradation signal for PKR remains unclear.

### PKR is actively depleted early in infection

We investigated when proteasomal degradation of PKR occurs. Early viral proteins are expressed prior to viral DNA replication, which is then followed by late viral protein expression. First, we examined the kinetics of PKR degradation to determine whether an early or late viral protein was likely responsible. We mock infected and infected CMT93 cells with MAV-1 at an MOI of 10 and collected lysates every six hours for 24 hours and analyzed them by immunoblot with antibodies to PKR or MAV-1 early region 1A (E1A) protein, the first viral protein made during infection (68). In infected cells, PKR degradation was first detected at 12 hpi (Fig. 7A), and quantitation of five independent experiments showed that ∼20% of the starting levels of PKR protein remained at 24 hpi (Fig 7B).

**Figure 7.**
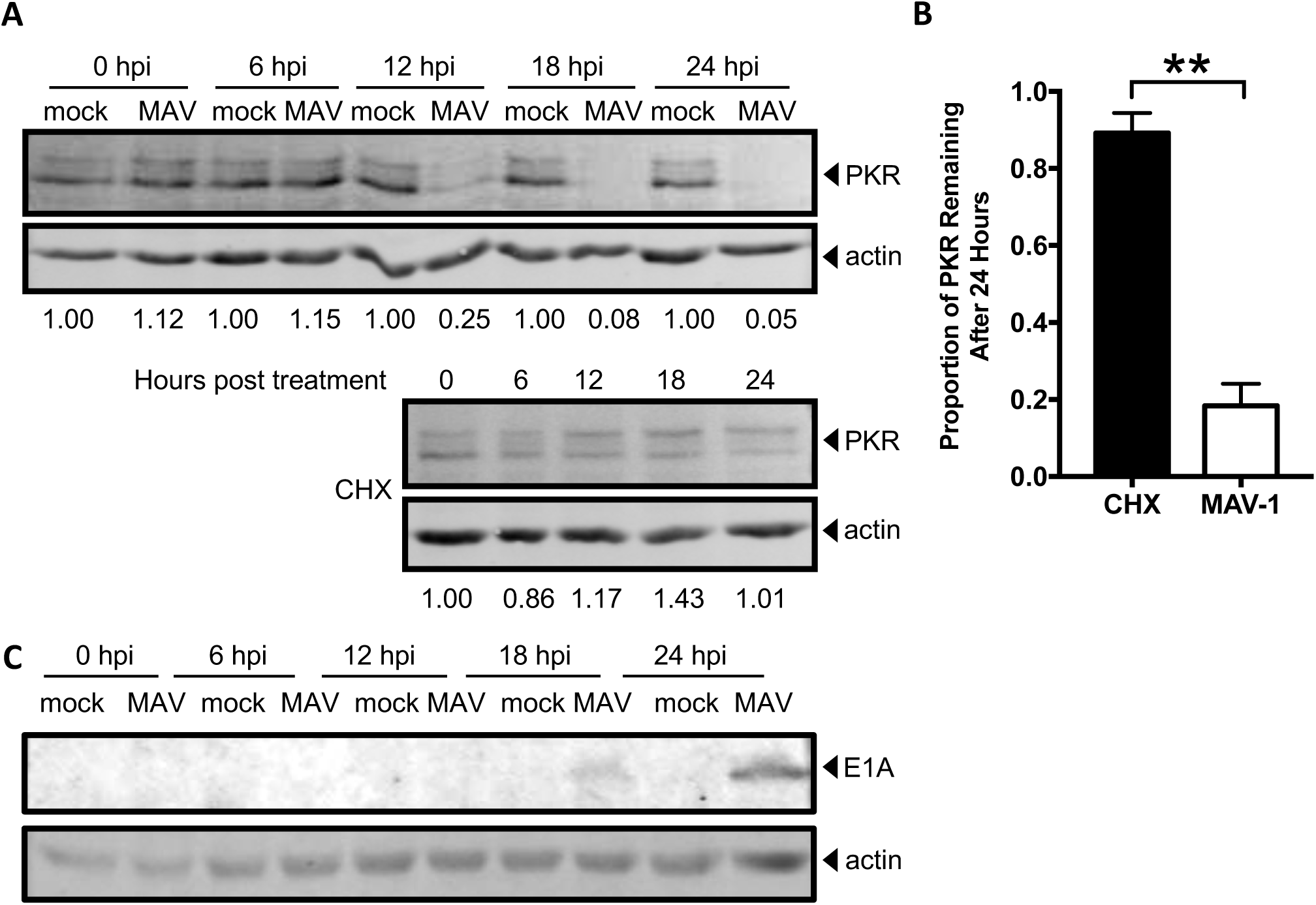
PKR is actively depleted early in infection. (A) CMT93 cells were infected with MAV-1 (MAV) at an MOI of 10 or mock infected (mock) (top), or uninfected cells were treated with 50 µg/mL cycloheximide (CHX, bottom) to inhibit elongation of protein synthesis. Cell lysates were analyzed by immunoblot with antibodies for PKR (D-20) and actin. Blots are representative of five independent experiments. (B) Densitometry quantitation of five independent experiments, \*\**P* ≤ 0.01. (C) CMT93 cell lysates from A were analyzed with a second immunoblot with antibodies for E1A and actin. Blots are representative of four replicates from two independent experiments. The E1A blot image was uniformly adjusted to a brightness of 30 and a contrast of 5 in Adobe Photoshop.

In parallel, to determine the half-life of PKR in uninfected CMT93 cells, we treated CMT93 cells with cycloheximide to halt protein translation and thus production of new PKR. We collected lysates every six hours for 24 hours and analyzed by immunoblot with antibodies to PKR. After 24 hours of cycloheximide treatment, approximately 90% of the starting levels of PKR protein remained (Fig. 7A bottom, and B). Comparing the results from MAV-1 infection (Fig. 7A top, and 7B) and cycloheximide treatment of uninfected cells (Fig. 7A bottom, and 7B), we conclude that MAV-1 was actively depleting PKR protein early in infection. E1A was detected by immunoblot at 18 hpi (Fig. 7C), whereas viral DNA replication was first detected at 24 hpi in CMT93 cells (Fig. S6). Thus the 18 hpi timepoint is considered an early time point during MAV-1 infection of CMT93 cells, prior to DNA replication, suggesting the involvement of an early viral protein in PKR depletion.

### An early viral function is required for PKR depletion by MAV-1

To determine whether viral gene expression or DNA replication are required for PKR degradation during infection, we infected C57BL/6 MEFs and CMT93 cells with UV-inactivated MAV-1 (which does not replicate viral DNA, Fig. S7). We infected cells at an MOI of 10 with WT MAV-1 or UV-inactivated MAV-1 and analyzed lysates from 24 and 48 hpi by immunoblot for PKR protein levels. In both cell types, while PKR was degraded by 24 hpi in the cells infected with WT MAV-1, PKR protein levels were unaffected at both time points in cells infected with UV-inactivated MAV-1 (Fig. 8A). This suggested that either gene expression or DNA replication was required for PKR degradation during MAV-1 infection.

**Figure 8.**
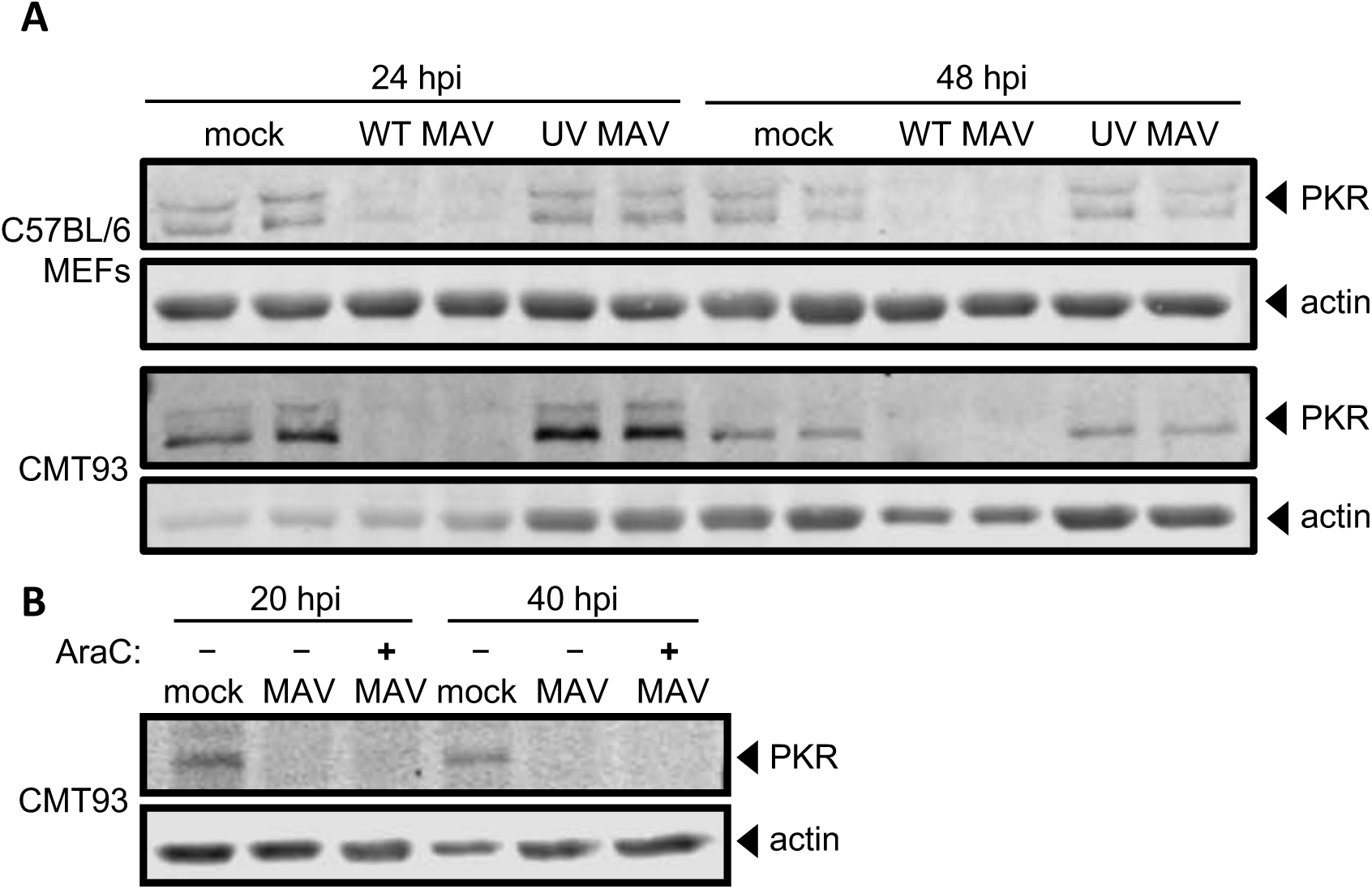
Early gene expression is required for PKR depletion by MAV-1. (A) Cells (as indicated at left) were infected with WT MAV-1 (WT MAV) or UV-inactivated MAV-1 (UV MAV) at an MOI of 10, or mock infected (mock). Cell lysates were analyzed by immunoblot with antibodies for PKR (D-20) and actin. Two independent wells were infected for each condition at both time points. (B) CMT93 cells were infected with WT MAV-1 (MAV) at an MOI of 10 or mock infected (mock). Infected cells were also treated (+) or not (-) with 20 µg/mL cytosine arabinasine (AraC), an inhibitor of DNA synthesis. Cell lysates were analyzed with antibodies for PKR and actin.

We addressed whether viral DNA replication is needed for PKR degradation. We mock infected or infected CMT93 cells with MAV-1 at an MOI of 10 and treated them with cytosine arabinoside (araC) at the time of infection to inhibit DNA synthesis (69, 70). This would allow the virus to infect the cell and produce early viral proteins, but would inhibit viral DNA replication and prevent late protein synthesis. We collected lysates at 20 and 40 hpi and analyzed them by immunoblot. We confirmed that araC treatment resulted in no late protein synthesis by performing an immunoblot for late virion proteins (Fig. S8). In samples treated with araC, PKR degradation was seen at 20 and 40 hpi (Fig. 8B), indicating that DNA replication was not required for PKR degradation. Taken together, the results of Fig. 8 are consistent with early viral gene expression prior to DNA replication being involved in induction of PKR degradation by MAV-1.

## Discussion

We have demonstrated here that PKR is antiviral in MAV-1 infections of cultured cells. Surprisingly, MAV-1 infection of primary and established cultured cells depleted PKR. The depletion was not due to reduced steady-state levels or reduced translation of PKR mRNA. Instead, we showed that PKR depletion is inhibited by proteasome inhibitors, implicating proteasomal degradation of PKR. Several lines of evidence suggest that the degradation is due to a viral early function.

PKR is an IFN-inducible gene product that is an important component of the innate immune response (1, 49). However, not all viruses have increased virulence in PKR^−/−^ MEFs, including EMCV and vaccinia virus (51, 71). While hAds produce VA RNAs that inhibit PKR antiviral activity during infection (10, 72), MAV-1 does not produce such VA RNAs, and how MAV-1 infection is affected by PKR is first described in this report. When we infected PKR^−/−^ MEFs with MAV-1, viral DNA yields were 5 to 6 times higher than viral DNA yields from wild type MEFs (Fig. 1), indicating that PKR plays an antiviral role during MAV-1 infection. At 48 hpi, the viral DNA yield from the N-MEFs was nearly 4 times higher than the C-MEFs, but by 72 hpi the viral DNA yields from both types of PKR^−/−^ MEFs were similar to each other and significantly increased compared to wild type MEFs (Fig. 1). This difference in viral replication kinetics between the two types of PKR^−/−^ MEFs may be due to differences in expression level and activity of the PKR fragments reportedly produced by them; we have not assayed PKR fragment production in our cells.

We examined PKR protein levels during MAV-1 infection and found that PKR was depleted from the cells as early as 12 hpi (Fig. 7). Depletion was seen in a wide variety of cell types, including immortalized C57BL/6 MEFs, primary C57BL/6 peritoneal macrophages, and CMT93 mouse colon carcinoma cells. Once depleted, PKR protein levels never returned to mock infected PKR protein levels during infection. Activation (transautophosphorylation) of PKR (6-9) was not required for this depletion, because kinase-dead mouse PKR was also depleted from K271R SV40-MEFs during infection (Fig. 2C). Both PKR and phospho-PKR were depleted in all cell types examined.

We examined several possibilities that could explain depletion of PKR protein, including PKR mRNA levels and alterations in translation. PKR mRNA levels remained unchanged during MAV-1 infection in C57BL/6 MEFs at 24 hpi and were increased in primary C57BL/6 peritoneal macrophages during MAV-1 infection (Fig. 3), times when the PKR protein levels were depleted (Fig. 2A). While PKR mRNAs in C57BL/6 MEFs were depleted 33% at 48 hpi and 40% at 72 hpi compared to mock lysates, this is not sufficient to explain the 84% and 94% reduction, respectively, in PKR protein levels at those time points (Fig. 2B). More likely, the reduction in PKR steady-state mRNA levels at the late infection time points can be attributed to other effects from viral infection, including the degradation or inhibition of proteins that induce PKR expression. For example, p53 is capable of binding to the PKR promoter and inducing its expression (73), but MAV-1 proteins cause p53 proteolysis (66). Together our results in C57BL/6 MEFs and macrophages suggest the virus does not cause PKR protein depletion by reducing PKR steady-state mRNA levels.

The differences in total PKR mRNA levels during infection between C57BL/6 MEFs and primary peritoneal macrophages (Fig. 3) is possibly due to the fact that macrophages are an immune cell, while MEFs are not. PKR protein took almost 3 times as long to be completely degraded during infection in macrophages compared to MEFs (72 hours versus 24 hours) (Fig. 2A). Total PKR mRNA levels were 2 to 3 times higher in MAV-1-infected macrophages compared to mock infected macrophages, unlike the MEFs where total PKR mRNA levels were unchanged or reduced 33-40% during MAV-1-infection compared to mock infected MEFs. Since PKR is an IFN-stimulated gene (1, 2), higher levels of total PKR mRNA seen during infection in the macrophages suggests IFN induction. This suggests that the immune response mounted by the macrophages was greater than the immune response in the MEFs, and could help explain why PKR took longer to degrade in macrophages compared to MEFs.

We considered whether reduced PKR levels were due to reduced PKR protein translation. There was no change in the total amount of PKR mRNA bound to ribosomes during infection compared to uninfected cells, nor was there a significant change in the amount of actively translating PKR mRNA during infection (Fig. 4). We also found that the 5’ UTR of mouse PKR placed upstream of a reporter gene produced the same amount of reporter with and without MAV-1 infection. These data indicate that there are not translational effects of MAV-1 infection on PKR protein levels that could explain the depleted PKR levels we observed.

Inhibiting lysosomal degradation resulted in no change in PKR depletion in infected cells (Fig. 5A), but adding proteasome inhibitors preserved PKR protein within cells (Fig. 6A and B). This indicates that PKR is not degraded by lysosomal degradation during viral infection, but by proteasomal degradation. Though PKR degradation was due to proteasome activity during MAV-1 infection, we were unable to demonstrate PKR ubiquitination, although we did detect ubiquitination of mouse p53 (Fig. S5). This inability to demonstrate PKR ubiquitination could be explained if at any given moment there were only low levels of ubiquitinated PKR present in the cell. Perhaps, increasing the time under MG132 treatment could increase the amounts of ubiquitinated proteins enough so that PKR ubiquitination could be seen. However, our inability to detect ubiquitinated PKR is consistent with a similar inability to identify PKR ubiquitination by RVFV NSs, even though NSs is known to recruit an E3 ligase to PKR (32). Alternatively, it is possible that in MAV-1 infection, PKR is degraded in a ubiquitin-independent manner, possibly because of intrinsic disordered regions of PKR or binding of regulating proteins to PKR that target proteins to the proteasome (74, 75).

Our experiments indicate that MAV-1 actively depletes PKR early in infection. Ongoing experiments are focused on determining the MAV-1 early protein(s) responsible for PKR degradation. Two possibilities are E4 proteins, the homologs of hAd E4orf6 and E4orf3, which we originally termed E4orfa/b and E4orfa/c, respectively (76). In human adenovirus, E4orf6 interacts with another early hAd protein, E1B 55K, to participate in an E3 ligase complex that ubiquitinates and degrades p53 via proteasomal degradation (77, 78). When MAV-1 E4orf6, E1B 55K, and mouse p53 are introduced by transfection into human cells, all three proteins interact and mouse p53 is degraded (66). If MAV-1 E4orf6 and E1B 55K form a similar complex in mouse cells, it may also degrade PKR. We have preliminary evidence that mouse p53 is ubiquitinated in C57BL/6 MEFs during MAV-1 infection, which suggests that the mouse p53 degradation seen in human cells could be paralleled by degradation of endogenous mouse p53 and mouse PKR in mouse cells, mediated by MAV-1 E4orf6 and E1B 55K during infection. Another hAd E4 protein, E4orf3, causes proteasomal degradation of transcriptional intermediary factor 1γ (79) and general transcription factor II-I (80) in a manner independent of hAd E4orf6 and E1B 55K. E4orf3 has SUMO E3 ligase and E4 elongase activity and induces sumoylation of general transcription factor II-I, leading to its proteasome-dependent degradation (80). MAV-1 E4orf3 may similarly have sumoylation activity that results ultimately in proteasome-dependent PKR degradation. Another possibility of a viral protein involved in PKR degradation is the protease encoded by MAV-1. The hAd protease is encapsidated in virions and proteolytically processes viral proteins IIIa, VI, VII, VIII, mu, and TP (81-84). However, we think it is unlikely that the MAV-1 protease degrades PKR, because we showed that UV-inactivated virus was unable to degrade PKR. We assume that UV treatment would not destroy the MAV-1 protease activity, just as HSV-1 VP16 activity is not altered by UV-inactivation of HSV-1 (85), but we have not tested this directly.

In summary, we demonstrated that PKR has an antiviral role during MAV-1 infection *in vitro*, because when PKR is mutated, viral replication in MEFs is significantly higher compared to wild type MEFs. Analysis of global PKR steady-state protein levels during infection showed complete PKR depletion by 72 hpi in multiple cell types, including immortalized and primary cells, with even faster kinetics in some. PKR transcription and translation were not decreased by MAV-1 infection, whereas proteasomal inhibition prevented PKR degradation. Taken together, these data suggest that MAV-1 causes PKR to be proteasomally degraded at a post-translational level. This work provides new insight into possible mechanisms of adenovirus inhibition of PKR by DNA viruses. PKR degradation may be induced by other adenoviruses that do not produce VA RNA, which includes all animal adenoviruses except primate adenoviruses and one type of fowl adenovirus (86).

## Materials and Methods

### Cells, virus, and infections

CMT93 cells (CCL-223) and C57BL/6 MEFs (SCRC-1008) were obtained from the American Type Culture Collection and passaged in Dulbecco’s modified eagle media (DMEM) containing 5% or 10% heat-inactivated fetal bovine serum (FBS), respectively, before use. Primary peritoneal macrophages were obtained from 6-10 week old C57BL/6 mice purchased from Jackson Laboratory (#000664) as described (87). Briefly, 6-10 week old C57BL/6 mice were injected intraperitoneally with 1.2 mL 3% thioglycolate and euthanized 3-5 days later. The abdominal skin was carefully removed, exposing the peritoneum, which was then injected with 5 mL of sterile phosphate-buffered saline (PBS). The abdomen was massaged gently, then the PBS containing the peritoneal macrophages was carefully withdrawn. The macrophages were centrifuged at 100 × g for 4 minutes, red blood cells lysed in lysis buffer (0.15 M ammonium chloride, 1 mM potassium bicarbonate, and 0.1 mM EDTA disodium salt) for 2 minutes at room temperature, centrifuged at 100 × g for 4 minutes, washed twice in PBS, resuspended in DMEM + 5% heat-inactivated FBS, and plated in 6 well plates. WT and PKR^−/−^ MEFs (termed PKR WT MEFs and N-PKR^−/−^ MEFs, respectively, throughout this paper) were obtained from Robert Silverman, Cleveland Clinic (88) and were passaged in DMEM containing 10% heat-inactivated FBS before use. PKR^−/−^ MEFs stably transfected with empty vector (termed C-PKR^−/−^ MEFs throughout this paper) were obtained from Dr. Gokhan Hotamisligil, Harvard University (89) and were passaged in DMEM containing 10% heat-inactivated FBS before use. WT (SV40-MEFs) and K271R PKR mutant (K271R SV40-MEFs) MEFs were obtained from Anthony Sadler, Hudson Institute of Medical Research (55) and were passaged in DMEM containing 10% heat-inactivated FBS before use.

Wild type mouse adenovirus type 1 (MAV-1) stock was prepared and titrated on mouse NIH 3T6 fibroblasts as described previously (90). WT MAV-1 was UV-inactivated by UV-treating 200 µL of virus for 10 min at 800 mJ/cm^2^. UV inactivation was confirmed by qPCR and plaque assay.

For infections, media was removed from cells and adsorption was performed in 0.4 mL of inocula for 6-well plate 35-mm wells (unless otherwise noted) for 1 hour at 37°C at the indicated MOIs (PFU/cell). After 60 minutes, 2 mL of DMEM+5% FBS was added without removing inocula; this time was designated as 0 hpi. For araC experiments, 20 µg/mL araC (Sigma C1768) was added at 0 hpi and replenished every 12-16 hours.

### Immunoblots

At room temperature, cells were washed once with PBS and Pierce™ RIPA lysis buffer (Thermo Scientific #89900) with 1x protease inhibitors (Protease Inhibitor Cocktail Kit, Thermo Scientific #78410) was added to the plate. The cells were allowed to lyse at room temperature for 10 minutes before being harvested and centrifuged at 4°C at 14,000 × g for 10 minutes to remove debris. Equivalent amounts of protein, determined by a BCA assay (Pierce BCA Protein Assay Kit, Thermo Scientific #23227), were acetone precipitated by incubating with 4x volume ice cold acetone overnight at −20°C. Precipitated proteins were pelleted at 4°C at 13,000 × g for 10 minutes and the pellets were dried for 30 minutes at room temperature. Pellets were resuspended in 10 µL Pierce™ RIPA lysis buffer (Thermo Scientific #89900), 3.25 µL NuPAGE 4x LDS Sample Buffer (Invitrogen Cat #NP0007), and 1.25 µL 1M DTT. Samples were incubated at 37°C for 10 minutes and then loaded into a well of an 8% acrylamide gel (8.3 cm wide × 7.3 cm high × 0.1 cm thick) with a 2.5% stacking gel, electrophoresed for 30 minutes at 50 V and 85 minutes at 150 V, and then transferred to a PVDF membrane (BioRad #1620177) for 1 hour at 100V at 4°C. Blots were blocked in 5% bovine serum albumin (BSA, Sigma A7906) in tris-buffered saline (BioRad #1706435) and 0.1% Tween 20 (Sigma P1379). Blots were probed with primary antibodies to detect mouse PKR (Santa Cruz D-20 sc-708, 1:2000, or B-10 sc-6282, 1:200), mouse actin (Santa Cruz sc-1616-R, 1:1000), MAV-1 E1A (AKO-7-147, 1:1000, described previously (68)), or MAV-1 late viral proteins (AKO 1-103, 1:1000, described previously (91, 92)). Secondary antibodies used were IRDye 800CW anti-rabbit (Li-Cor 925-32213, 1:15,000) or IgG peroxidase-conjugated anti-mouse (Jackson Immuno 515-035-062, 1:20,000). Blots were visualized by LI-COR Odyssey imaging (LI-COR Biosciences) or enhanced chemiluminescent substrates (Pierce ECL Western Blotting Substrate #32106) and X-ray film (Dot Scientific #BDB810). Densitometric quantification was performed on .tif files using ImageJ software from NIH (93).

To attempt to demonstrate PKR ubiquitination status during MAV-1 infection, C57BL/6 MEFs were transfected with GFP- or HA-epitope tagged ubiquitin plasmids (Addgene #11928 and #18712, respectively) 24 hours before infection. We used Polyplus jetPRIME transfection reagent (Polyplus #114-15) with 10 µg plasmid DNA and 30 µL jetPRIME reagent per 10 cm plate. At 12 hpi (36 hours post transfection), we treated mock and infected C57BL/6 MEFs with 10 µM MG132 (Sigma M7449) for 6 hours before collecting lysates at 18 hpi in HCN buffer (50 mM HEPES, 150 mM NaCl, 2 mM CaCl_2,_ 1% Triton X-100 (Sigma T9284), 1x protease inhibitors (Protease Inhibitor Cocktail Kit, Thermo Scientific #78410), and 5 mM N-ethylmalemide). The lysates were split into two aliquots, and 3 µg PKR (D-20 sc-708, discontinued) or 3 µg isotype rabbit polyclonal antibody (Jackson Immuno #011-000-002) was added to lysates. After rocking samples overnight at 4°C, 20 µL protein A agarose suspension (Calbiochem/Millipore #IP02-1.5ML) was added to each and samples were rocked at 4°C for 2 hours. After incubation, agarose was washed 3 times with 1 mL HCN buffer, resuspended in 40 µL 2x Laemmli buffer (BioRad #161-0737) with 5% 2-mercaptoethanol (Sigma M6250), and boiled for 10 minutes. Lysate supernatants remaining after the initial PKR immunoprecipitation were then immunoprecipitated again using the same procedure but with 4 µg anti-p53 (DO-1, Santa Cruz sc-126) or 4 µg isotype mouse monoclonal antibodies (ThermoFisher Scientific #02-6200). Immunoprecipitated proteins were immunoblotted for GFP- or HA-epitope tagged ubiquitin with antibodies for GFP (1:3,000, Roche #11814460001) or HA (1:4,000, Abcam #ab9110). No ubiquitin signal was detected from the PKR immunoprecipitations, though the positive control, p53, showed ubiquitin signal with both epitope-tagged ubiquitins (data not shown). Blots were also probed for PKR (1:200, PKR B-10 sc-6282), p53 (1:200, anti-p53 sc-98), and IRDye 800CW anti-mouse (Li-Cor 925-32212, 1:15,000) to confirm the immunoprecipitations were successful, and signals for both proteins were detected (data not shown).

### Viral DNA yield analysis by qPCR

Cells were washed twice with room temperature PBS and harvested by scraping into PBS, centrifuging at 100 × g for 4 minutes at 4°C, and resuspending in PBS. Total cellular DNA was purified using the Invitrogen PureLink DNA Purification Kit (Thermo Scientific #K1820-02) and quantitated by a NanoDrop Spectrophotometer. 10 ng of total cellular DNA was analyzed by qPCR using custom primers specific to MAV-1 E1A (mE1Agenomic Fwd: 5’ GCA CTC CAT GGC AGG ATT CT 3’ and mE1Agenomic Rev 5’ GGT CGA AGC AGA CGG TTC TTC 3’) and the results were normalized to GAPDH, which was analyzed using a GAPDH-specific primer/probe set (ThermoFisher Scientific, Mm99999915_g1, #4331182).

### mRNA analysis by qPCR

Cells were harvested by scraping into media, centrifuging at 100 × g for 4 min at 4°C, and washing the cell pellet three times with ice-cold PBS. RNA was purified using the Qiagen RNeasy Mini Kit (Qiagen #74134) and stored at −80°C. 125 ng of RNA per sample was used to make cDNA using the High Capacity cDNA Reverse Transcription Kit (Applied Biosystems #4368814), and 2 µL of the cDNA was analyzed by qPCR using a primer/probe set specific to mouse PKR sequence (Thermo Fisher, Mm01235643_m1, #4331182). The results were normalized to GAPDH, which was analyzed using a GAPDH-specific primer/probe set (ThermoFisher Scientific, Mm99999915_g1, #4331182). Arbitrary units were calculated as follows: Mean C_T_ PKR – mean C_T_ GAPDH = ΔC_T_ for sample. Arbitrary unit = 2^-ΔC_T_.

### Proteasome inhibition

C57BL/6 MEFs were infected at an MOI of 10, and DMSO, 1 µM MG132 (Sigma M7449), or 1 µM bortezomib (Selleckchem #S1013) were added to the media after a 1 hour adsorption. At 24 hpi, cells were washed once with room temperature PBS, and Pierce™ RIPA lysis buffer (Thermo Scientific #89900) with 1x protease inhibitors (Protease Inhibitor Cocktail Kit, Thermo Scientific #78410) was added to the plate. The cells were allowed to lyse at room temperature for 10 minutes before being harvested and centrifuged at 4°C at 14,000 × g for 10 minutes to remove debris.

### Lysosome inhibition and DQ BSA assay

CMT93 cells were infected at an MOI of 10. After a 1 hour adsorption, 10 µL water, 10 mM NH_4_Cl (final concentration, Baker Chemical Company #0660-1), or 60 µM chloroquine (final concentration, Sigma C6628) was added to the media. At 24 hpi, at room temperature, cells were washed once with PBS and Pierce™ RIPA lysis buffer (Thermo Scientific #89900) with 1x protease inhibitors (Protease Inhibitor Cocktail Kit, Thermo Scientific #78410) was added to the plate. The cells were allowed to lyse at room temperature for 10 minutes before being harvested and centrifuged at 4°C at 14,000 × g for 10 minutes to remove debris.

The DQ BSA assay was performed as described (60). Briefly, C57BL/6 MEFs and CMT93 cells were plated at 1.5 × 10^5^ cells/plate or 3 × 10^5^ cells/plate, respectively, in MatTek Glass Bottom Microwell Dishes (Part No: P35G-1.5-14C) with 2 mL of DMEM + 10% or 5% FBS, respectively. The next day, the cell media was treated with 10 µL water, 10 mM NH_4_Cl (final concentration), or 60 µM chloroquine (final concentration, Sigma C6628). Four hours after adding inhibitors, DQ Red BSA (Invitrogen Cat #D12051) was added to the media to a final concentration of 5 µg/mL in 2 mL DMEM + 10% or 5% FBS, respectively. At 24 hours post treatment, the cells were imaged on a Nikon TE300 inverted microscope equipped with a mercury arc lamp, Plan-Apochromat 60x, 1.4 NA objective, cooled digital CCD camera (Quantix Photometrics, Tucson, AZ), and a temperature-controlled stage, set at 37°C. To image the DQ-BSA, we used an excitation filter centered at 572 nm and an emission filter centered at 635 nm. The exposure time was the same for all images.

### Ribosome pelleting

Ribosomes were pelleted as described (94). Briefly, C57BL/6 MEFs were plated on 10 cm plates, 3×10^5^ cells per plate. The next day, the cells (∼90% confluent) were infected with MAV-1 at an MOI of 5. C57BL/6 MEF lysates were collected at 48 hpi by scraping the cells in ice-cold PBS containing 100 µg/mL cycloheximide (Sigma C7698), pelleting, and resuspending in lysis buffer, which was 10 mM HEPES pH 7.5, 100 mM KCl, 5 mM MgCl_2_, 4 mM DTT, 0.5% NP-40, 100 µg/mL cycloheximide, 20 U/mL RNasin (Promega # N2511), 10% sucrose, and 1x protease inhibitors (Protease Inhibitor Cocktail Kit, Thermo Scientific #78410). Cells were lysed by passage through a chilled 26G needle five times and cleared by centrifugation for 10 min at 21,000 × g at 4°C. 400 µL of cleared lysate (10 OD_260nm_ units) was layered onto 25% sucrose and centrifuged 29,500 rpm in an SW41 rotor (107,458 rcf average) for 4 hours at 4°C. After pelleting, the supernatant was removed with a micropipet and 350 µL of 4°C Buffer RLT Plus (from Qiagen RNeasy Mini Kit) was added to the pellet to collect the RNA. RNA was purified immediately using the Qiagen RNeasy Mini Kit (Qiagen #74134) and stored at −80°C until analysis.

### Polyribosome Gradients

C57BL/6 MEFs were plated on 10 cm plates, 2 × 10^6^ cells per plate. The next day, the cells were infected with MAV-1 at an MOI of 2. Following a standard protocol (95), 5 minutes prior to collection, cycloheximide was added at a final concentration of 100 µg/mL and incubated at 37°C. Cells were collected at 24 hpi by scraping in ice-cold PBS containing 100 µg/mL cycloheximide, pelleting, and resuspending in 500 µL lysis buffer (20 mM Tris-Cl, 150 mM NaCl, 15 mM MgCl_2_, 8% glycerol, 20 IU/mL SUPERase•In (ThermoFisher Scientific Cat# AM2696), 80 IU/mL Murine RNAse Inhibitor (New England BioLabs Cat# M0314S), 0.1 mg/mL heparin (Sigma H3393-50), 0.1mg/mL cycloheximide, 1 mM DTT, 1x protease inhibitor (Protease Inhibitor Cocktail Kit, Thermo Scientific #78410), 20 IU/mL Turbo DNAse (ThermoFisher Scientific Cat# AM2238), and 1% Triton X-100 (Sigma T9284). Cells were lysed by passaging through a chilled 26G needle ten times, vortexing for 30 seconds, and then incubating on ice for 5 min. Lysates were cleared by centrifugation for 5 min at 14,000 × g at 4°C. 500 µL of cleared lysate (10 OD_260nm_) was layered onto a 10-50% sucrose gradient and centrifuged at 35,000 rpm in an SW41 rotor (151,000 rcf) for 3 hours at 4°C. After centrifugation, gradients were pumped out of the top with a Brandel BR-188 Density Gradient Fractionation System with a continuous reading of the OD_254 nm_. From 24-34 fractions (350-500 µL) were collected. RNA was purified from selected fractions immediately using the Qiagen RNeasy Mini Kit (Qiagen #74134) and stored at −80°C until analysis by RT-qPCR.

### Statistical analyses

Data were analyzed with GraphPad Prism 7.0 software. For qPCR and densitometry analyses, the data were analyzed by individual Mann-Whitney tests. A value of *P* < 0.5 was considered significant.

## Acknowledgments

We thank Robert Silverman, Gokhan Hotamisligil, and Anthony Sadler for PKR mutant and wild type cells. We thank Katelyn Green, Becky Billmire, and Peter Todd for the use of and assistance with the Brandel BR-188 Density Gradient Fractionation System and for providing the nanoluciferase and firefly luciferase plasmids. We thank Zachary Mendel and Joel Swanson for assistance with imaging the DQ BSA assay. We thank Jason Weinberg, Michael Imperiale, Christiane Wobus, Billy Tsai, Andrew Mehle, Mitch Ledwith, Mike Mathews, Chuck Samuel, and Britt Glaunsinger for helpful discussions. We thank Michael Imperiale and Christiane Wobus for comments on the manuscript.

This work was supported by HHS/NIH/National Institute of Allergy and Infectious Diseases (NIAID), 5 R01AI091721 and 1 R01AI133935 (Katherine R. Spindler); and NIH Cellular and Molecular Biology Training Grant T32 GM007315, and Rackham Pre-Candidate and Candidate Student Research Grants (Danielle E. Goodman).

## Supplemental Figure Legends

**Supplemental Figure 1.**
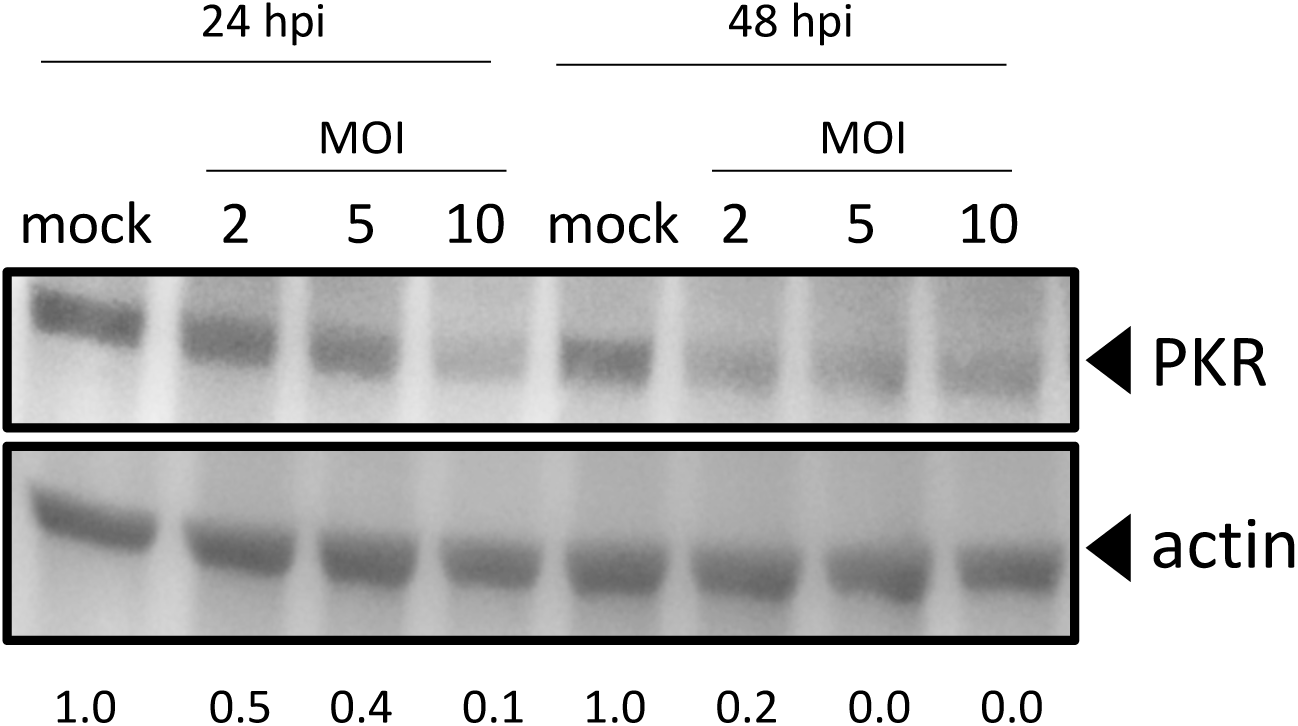
PKR is depleted at MOIs of 2, 5, and 10. C57BL/6 MEFs were infected with MAV-1 at an MOI of 2, 5, or 10 (2, 5, 10) or mock infected (mock). Cell lysates were analyzed by immunoblot with antibodies for PKR (B-10) and actin. Densitometry quantitation using ImageJ is listed below each lane; for each time point, the infected samples were normalized to the mock. The PKR blot image was uniformly adjusted to a brightness of 150 in Adobe Photoshop.

**Supplemental Figure 2.**
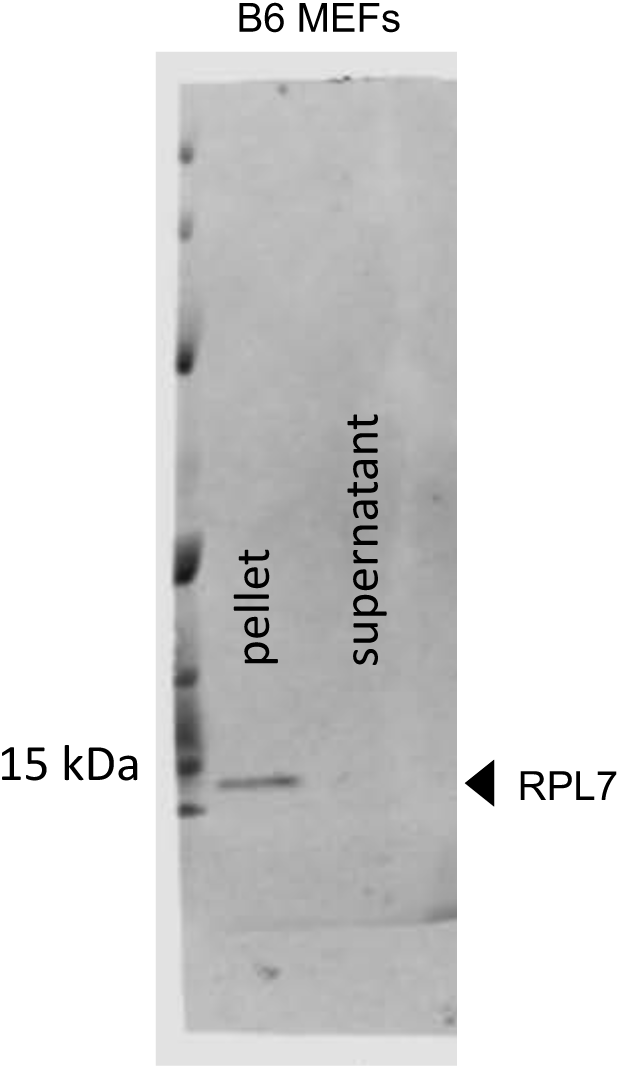
Ribosome pelleting. To confirm that most ribosomes ended up in the pellet after centrifugation through sucrose (Fig. 4A), a sample of the pellet and the corresponding supernatant were analyzed by immunoblot with antibodies for RPL7 (ribosomal protein L7, Abcam, 1:2000, ab72550).

**Supplemental Figure 3.**
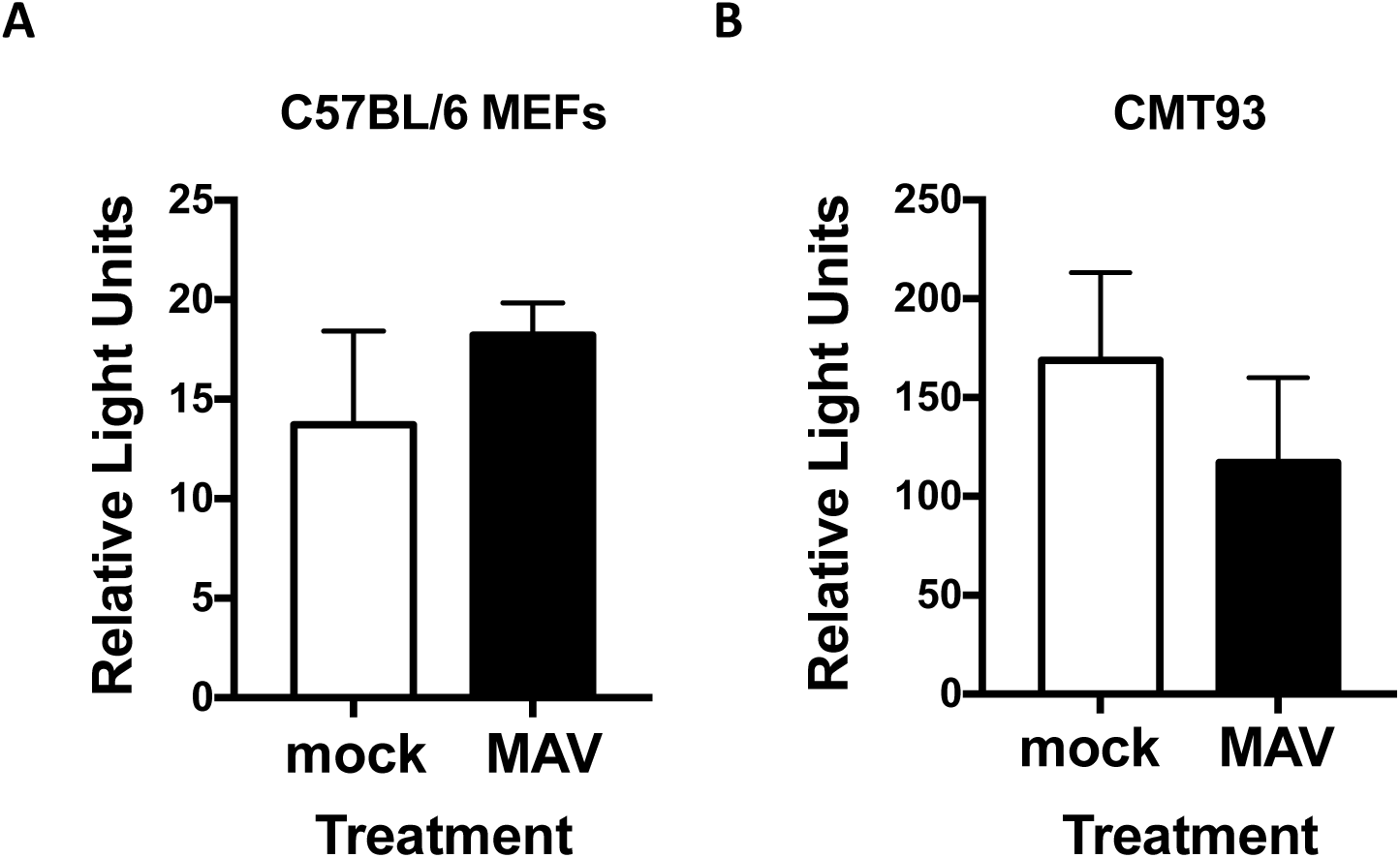
PKR mRNA 5’ UTR does not result in altered reporter protein levels upon MAV-1 infection. (A) C57BL/6 MEFs or (B) CMT93 cells were co-transfected with pmPKR5UTRfullNL or AUG-NL-3xFLAG and pGL4.13 using jetPRIME reagents (Polyplus #114-15) using the standard Polyplus protocol, with 200 ng total of plasmid and 300 µL of jetPRIME reagent per 35 mm well. At 24 hours after transfection, the cells were infected with MAV-1 at an MOI of 10. At 24 hpi, cells were lysed in 70 µL/well Glo Lysis Buffer (Promega Corp.). After lysing, 25 µL of each lysed sample and 25 µL of OneGlo or NanoGlo (Promega Corp.) was added to two wells in a black 96-well plate (Fisher Scientific #07-000-634). After 5 minutes, the plate was read on a Promega GloMax luminometer. Relative light units from the pmPKR5UTRfullNL plasmid were normalized to the firefly luciferase and positive control plasmids. Graphs are representative of 7-9 biological replicates per treatment group.

**Supplemental Figure 4.**
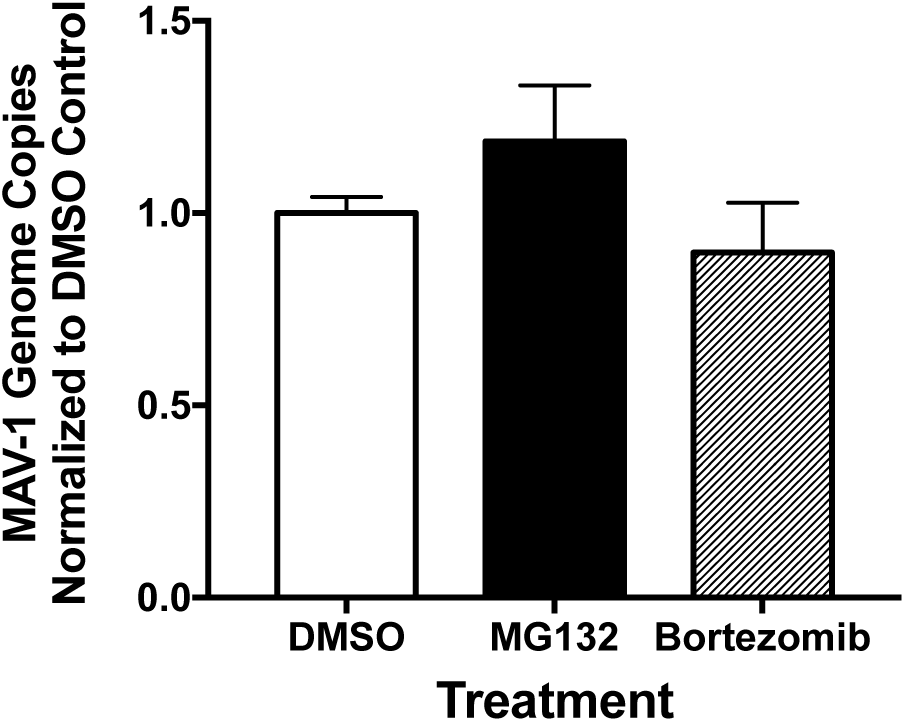
MG132 and bortezomib treatment do not affect MAV-1 replication at 24 hpi. C57BL/6 MEFs were infected with MAV-1 at an MOI of 10 and treated with DMSO (vehicle for inhibitors), 1 µM MG132 or bortezomib, and collected at 24 hpi. DNA was purified from cell pellets and analyzed for MAV-1 genome copies by qPCR. Graph is representative of five biological replicates per treatment group. Error bars are standard error of the mean (SEM). \**P* ≤ 0.05.

**Supplemental Figure 5.**
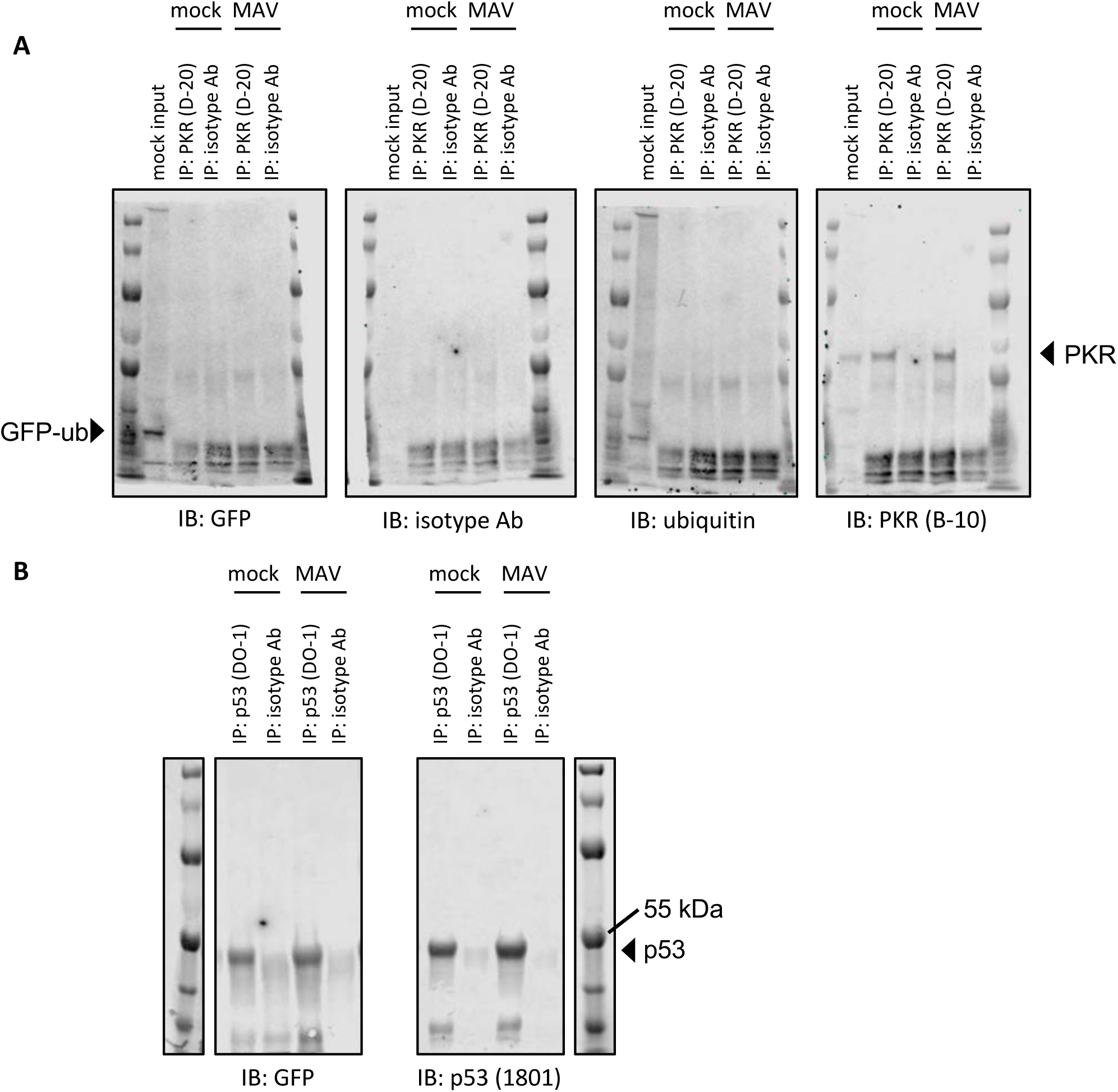
PKR is not detectably ubiquitinated during MAV-1 infection. CMT93 cells were transfected with a GFP-ubiquitin plasmid using standard Polyplus transfection protocols. At 24 hpt, the cells were infected with MAV-1 (MAV) at an MOI of 5 or mock infected (mock) and treated with 10 µM MG132 at 6 hpi. Cell lysates were collected at 12 hpi and immunoprecipitated with (A) PKR (D-20) or an isotype control antibody or (B) p53 (DO-1) or an isotype control antibody. Immunoprecipitated samples were analyzed by immunoblot with GFP, ubiquitin, isotype, PKR (B-10), or p53 (1801) antibodies. Input lane (mock input) contains 0.008 volume of mock infected lysate (relative to volume in immunoprecipitations). For (B) GFP and p53 were probed on separate duplicate blots.

**Supplemental Figure 6.**
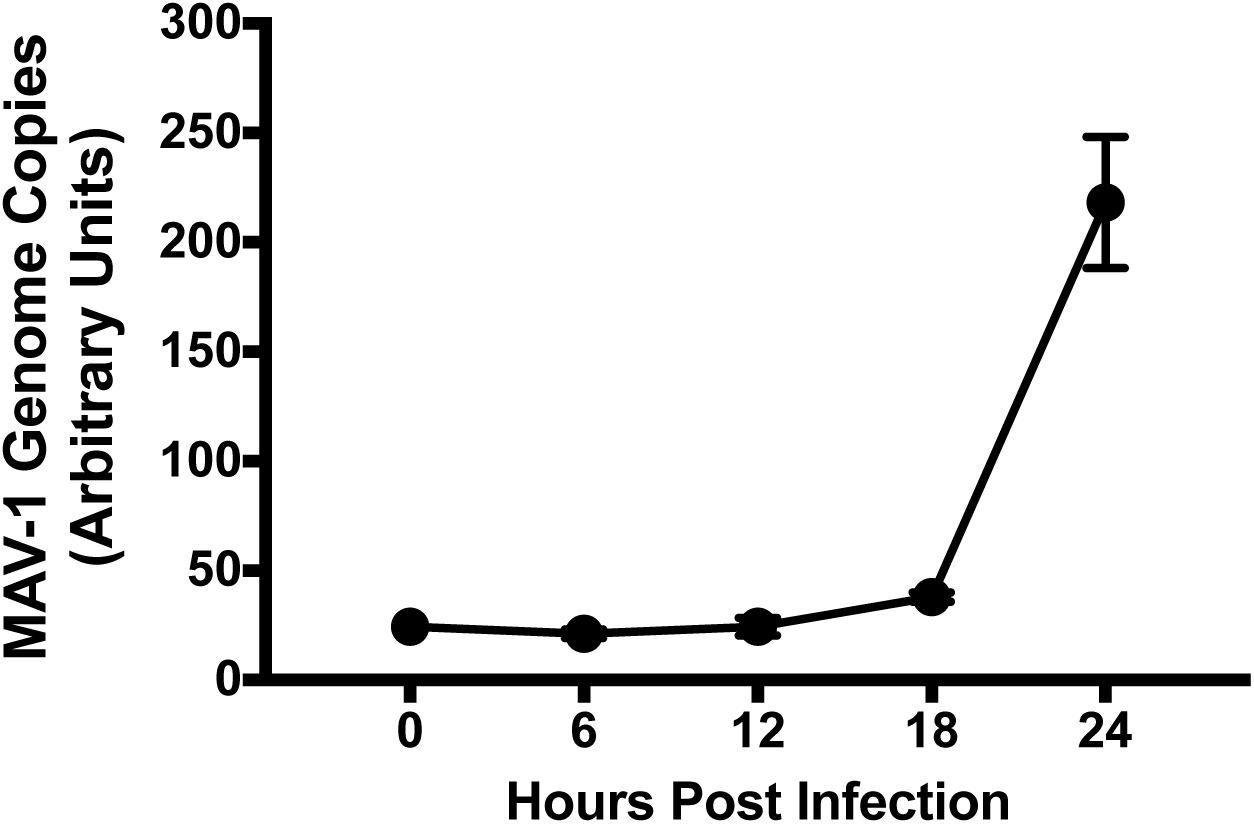
Viral DNA replication can be detected at 24 hpi by qPCR. CMT93 cells were infected with MAV-1 (MAV) at an MOI of 10 and collected every 6 hours for 24 hours. DNA was purified from cell pellets and analyzed for MAV-1 genome copies by qPCR. Graph is representative of four to five biological replicates per treatment group. Error bars are standard error of the mean (SEM).

**Supplemental Figure 7.**
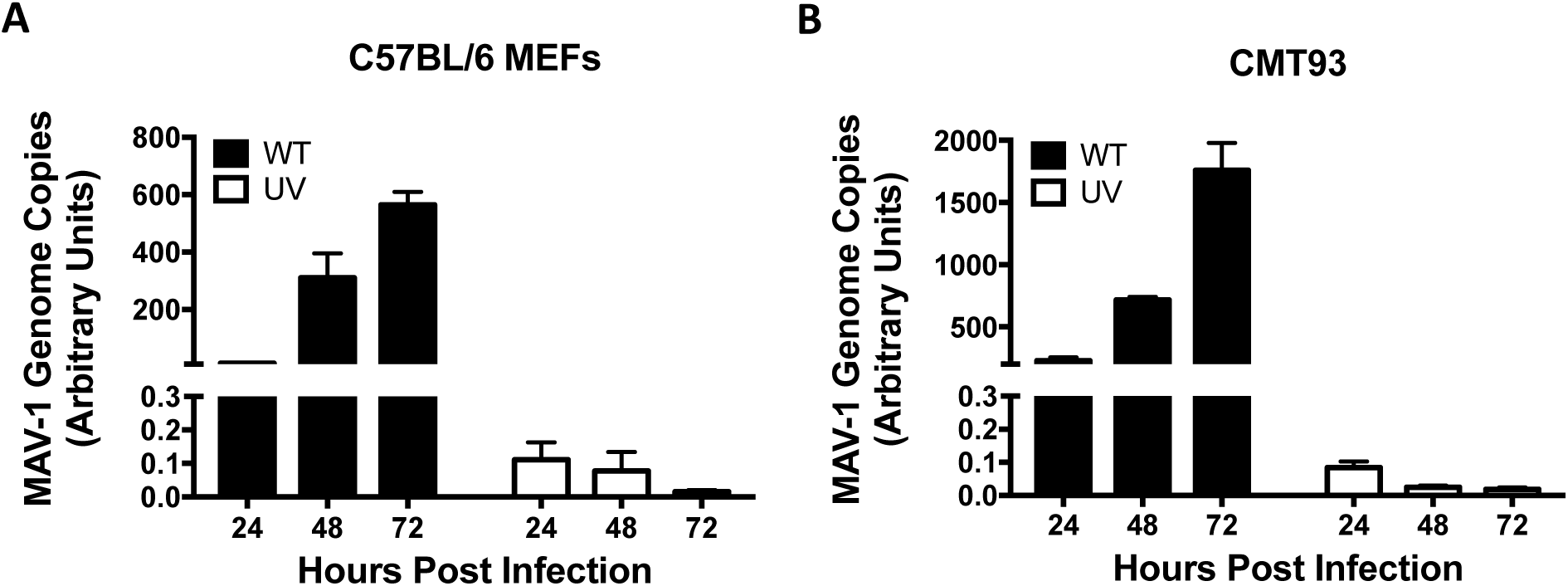
UV-inactivated virus does not replicate viral DNA. (A) C57BL/6 MEFs or (B) CMT93 cells were infected with WT MAV-1 (WT) or UV-inactivated MAV-1 (UV) at an MOI of 10 and collected at indicated times. DNA was purified from cell pellets and analyzed for MAV-1 genome copies by qPCR. Graphs are representative of three to four biological replicates per treatment group. Error bars are standard error of the mean (SEM).

**Supplemental Figure 8.**
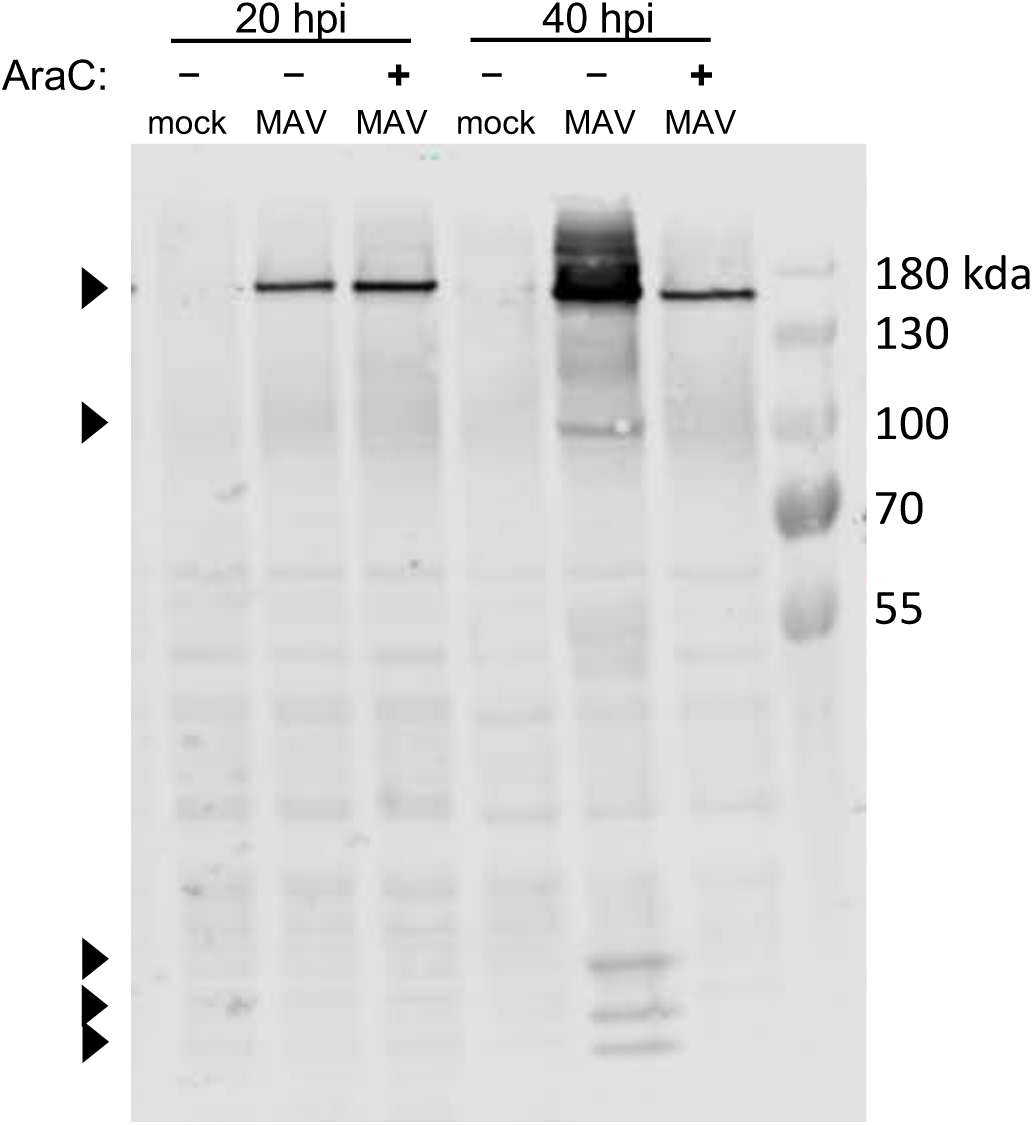
AraC treatment inhibited late protein expression. CMT93 cells were infected with WT MAV-1 (MAV) at an MOI of 10 or mock infected (mock). Infected cells were also treated (+) or not (-) with 20 µg/mL cytosine arabinasine (araC), an inhibitor of DNA synthesis. Cell lysates were analyzed with antibodies for late virion proteins (AKO1-103, 1:1000). Arrowheads indicate late viral proteins.

